# DeTOKI identifies and characterizes the dynamics of chromatin topologically associating domains in a single cell

**DOI:** 10.1101/2021.02.23.432401

**Authors:** Xiao Li, Zhihua Zhang

## Abstract

The human genome has a dynamic, well-organized hierarchical 3D architecture, including megabase-sized topologically associating domains (TAD). TADs are a key structure of the genome regulating nuclear processes, such as gene expression, DNA replication and damage repair. However, owing to a lack of proper computational tools, TADs have still not been systematically and reliably surveyed in single cells. In the present work, we developed a new algorithm to <u>de</u>code <u>T</u>AD b<u>o</u>undaries that <u>k</u>eep chromatin <u>i</u>nteraction insulated (deTOKI) from ultra-sparse Hi-C data. By nonnegative matrix factorization, this novel algorithm seeks out for regions that insulate the genome into blocks with minimal chance of clustering. We found that deTOKI outperformed competing tools and that it reliably identified TADs with single-cell Hi-C (scHi-C) data. By applying deTOKI, we found that domain structures are prevalent in single cells. Further, although domain structures are highly dynamic between cells, TADs adhere to the ensemble, suggesting tight regulation of single-cell TADs. Finally, we found that the insulation properties of TAD boundaries have major effect on the epigenetic landscape in individual cells. In sum, deTOKI serves as a powerful tool for profiling TADs in single cells.

## Introduction

The eukaryote genome in the nucleus is folded into a hierarchical configuration^1,2^ as revealed by imaging technologies^3^ and chromosome conformation capture (3C) based technologies^4–12^, e.g., Hi-C^8^ The hierarchical configuration consists of chromosomal territories^8,13,14^, A and B compartments^8^, domain structures, such as topologically associating domains (TADs)^14,15^, compartment domains^16^, or CTCF loop domains^17^, and chromatin loops^17–19^ Such configurations have been routinely discussed in many studies^2^. TADs might be the most investigated chromatin feature in the literature since their disruption can cause severe diseases^20^, including cancer^21^.

The TAD structures were mostly revealed by Hi-C in bulk cells^8^, while the existence and biogenesis of TADs in individual cells remain unclear. Super-resolution imaging data has shown the existence of and the variations in the TAD-like domain structures in single cells^22^. Given the large cell to cell variations of chromatin architecture observed in individual cells, TADs could be a partially emergent property of a cell population. That is, the dynamics of chromatin in single cells *per se* may generate, at least in part, the TADs we observed in the bulk cells^23–25^. Since the origin and dynamics of TADs are keys to understanding gene regulation^26,27^, unraveling the nature of single-cell TAD structure is essential. However, systematic survey of TAD structure and its dynamics in single cells remains a major challenge in the field.

A long list of TAD detection tools are available in the literature, and the methods are sophisticated and diverse^28,29^ TADs were first identified using certain local genomic or topologic features, e.g., the directionality index (DI)^14^, the insulation score (IS)^30^, the arrowhead score^18^, IC-Finder^31^ and ClusterTAD^32^. Later, methods based on probabilistic models with certain assumption about the data distributions were developed, such as GMAP^33^, PSYCHIC^34^, HiCseg^35^, TADbit^36^ and TADtree^37^. Some other tools utilize dynamic programming to optimize a global object function, e.g., Armatus^38^ and Matryoshka^39^. When treating the Hi-C matrix as a network connection matrix, an entire toolbox is available from graph theory, e.g., MrTADFinder, 3DnetMod^40,41^, and we recently developed deDoc^42^. However, comparisons have shown that almost none of them worked reliably with ultra-low resolution Hi-C data^28,43^ Among all TAD predictors, IS and deDoc worked the best with low resolution Hi-C^30,42^; however, TADs are virtually undetectable in experimental single-cell Hi-C data.

The inadequate handling of single-cell Hi-C data by current TAD prediction methods stems from the ultra-sparsity of chromatin interactions. A single cell has two copies for any given locus, and therefore, at most only two copies of Hi-C ligations could possibly exist in the single-cell Hi-C (scHi-C) libraries for that locus. Thus, the fluctuations from stochastic chromatin interactions *per se,* or from PCR proliferation, have exponential effect on the final scHi-C sequencing data. Consequently, to systematically survey the TAD structure, we need a computational tool that is able to reliably process such ultra-sparse data from scHi-C.

Non-negative matrix factorization (NMF) consists of a group of algorithms in multivariate analysis where a non-negative matrix is factorized into two or more non-negative matrices^44^. The NMF has been widely used in processing single-cell omics data, e.g., coupled NMF^45^ The advantage of NMF is its low rank representation, which retrieves key information embedded in the noisy sparse data. As a sparse non-negative matrix, the sparsity issue of scHi-C data can also be solved by NMF. Therefore, we developed a new method using NMF to <u>de</u>code <u>T</u>AD b<u>o</u>undaries that <u>k</u>eep chromatin <u>i</u>nteraction isolated (deTOKI) from ultra-sparse Hi-C data. We present evidences that deTOKI can reliably predict TAD domain structures at the single-cell level. TAD structures are highly dynamic between cells. However, by applying deTOKI to public experimental single-cell Hi-C data, we found that although the TAD structure is highly dynamic between the cells, the TADs adhere to the ensemble, suggesting their tight regulation. Finally, we found that the insulation property of the TAD boundaries also has a major effect on the epigenetic landscape in individual cells.

## Results

### A novel TAD detector (deTOKI) for ultra-low resolution Hi-C data

We developed a novel algorithm, named deTOKI, to detect TAD structures using using ultra-sparse Hi-C contact matrices. The deTOKI takes advantage of a key property of TADs. Namely, that its topology distribution is relatively consistent with respect to number and length between cell types^14^ Briefly, for any given genome segment, deTOKI applies non-negative matrix factorization (NMF) to decompose the Hi-C contact matrix into genome domains that may be spatially segregated in 3D space (Fig.1a). Nonnegative matrix factorization is an algorithm for decomposition of a nonnegative matrix into a product of two nonnegative matrices, in which the n_components represent the common dimensions between the two decomposed matrices. As the n_components normally are substantially smaller than the dimensions of the origin matrix, NMF is a commonly used algorithm to perform dimension reduction^46^. To speed up the algorithm, deTOKI divides the chromosomes into 8Mb windows, and the clusters in the middle 4Mb of each window are reported as the predictions (Method, Fig.1, Supplementary Fig.1 and 2). The alternative local optimal solutions in the structure ensemble are achieved by summarizing deTOKI’s predictions with multiple random initiations (see Methods).

**Fig. 1.**
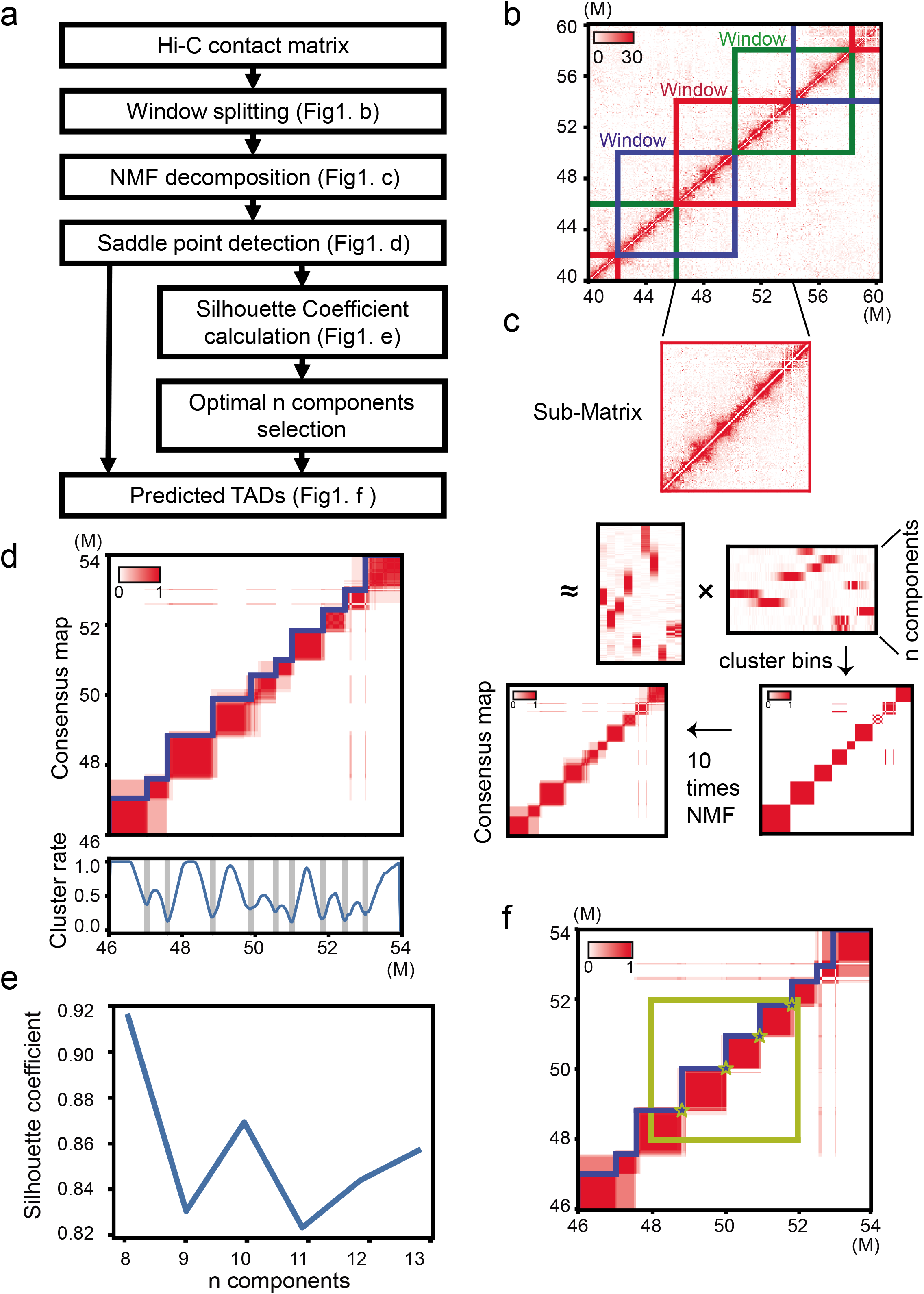
Workflow of deTOKI. **(a)** The flow chart of deTOKI. The genome is split into 8MB overlapping sliding windows. As an example, **(b-f)** shows the key steps for a window in chr19. The data are from IMR90 cells^14^. In **(b)**, each square in blue, red and green represents a sliding window. In (**c**), NMF was performed on the contact matrix of a window, and bins were clustered based on the factor matrix. After 10 rounds of NMF, a consensus map was generated **(d)**, in which the detected clustering change points are shown in sawtooth, and the blue curve in the bottom panel represents the clustering rate of each bin. **(e)** The blue curve represents the silhouette coefficient corresponding to each alternative value of “n_components”. (**f**) The yellow square highlighted the middle 4Mb region for which clustering change points (yellow stars) were reported as TAD boundaries.

To assess the deTOKI performance, we focused on two major characteristics of chromatin architecture at single cell level, i.e., the data sparsity and the cell to cell variations, and assessed them with down-sampled experimental data and with simulated data, respectively.

### deTOKI worked well in down-sampled bulk Hi-C at single-cell level

To assess the performance of deTOKI under the condition of sparse input, we mainly compared it with two publically available algorithms, e.g., Insulation score (IS)^30^ and deDoc^42^. These two method were chosen because they were judged to be the most robust methods with sparse data in our previous comprehensive assessments of TAD predictors^43^. In addition, we also compared it with two recently published algorithms that designed for sparse data, SpectralTAD^47^ and GRiNCH^48^ The sparsity was defined as the proportion of entries in the Hi-C matrix that have value zero after excluding the unmappable genome regions, e.g., centromeres, for a given chromosome. The assessment was done for all chromosomes in 40kb bins, and was down-sampled at the rate of 1/800 from the high resolution Hi-C data^14^. The down-sampled data set consisted of about 0.44M contacts, mimicking the sequencing depths of public scHi-C datasets, e.g., the median of the data generated by Flyamer and colleagues (hereafter termed the Flyamer’s data^49^) was 0.339M (Fig. 2a and 2b).

**Fig. 2.**
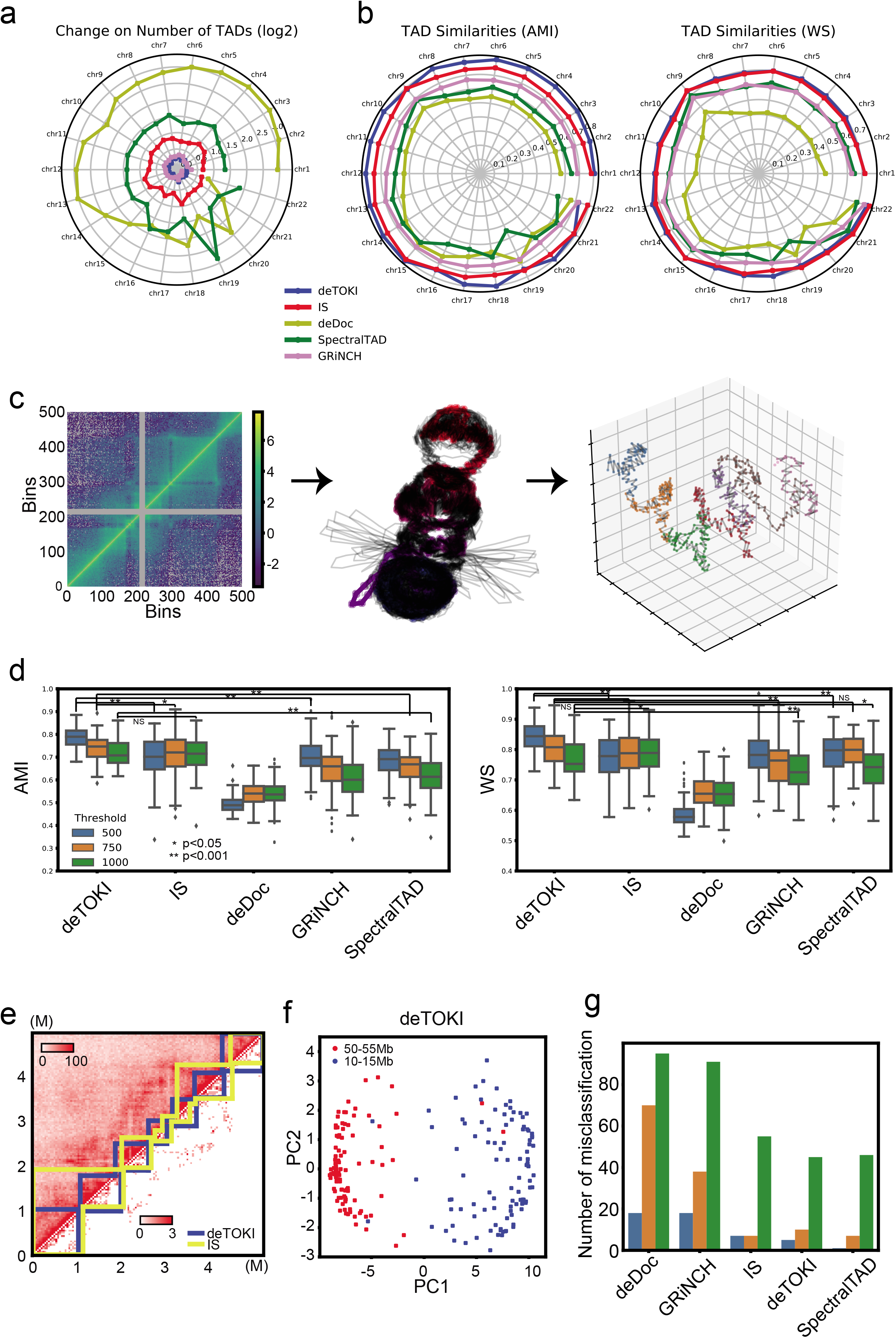
Comparison of TAD callers on simulated single-cell Hi-C data. The simulation is based on data form IM90^14^. Panels (a) and (b) shows the average results of 20 independent down-samplings in each chromosome. **(a)** The (log2) change in the number of predicted TADs. **(b)** The similarity of TADs, as inferred by AMI and WS, between the raw data and the down-sampled data. **(c)** Workflow of the single-cell Hi-C simulation. From left to right the panels represent the normalized Hi-C contact matrix of chr18:50-55Mb for GM12878 ensemble Hi-C from Rao’s data^18^, an ensemble of 100 modeled 3D structures of this region, and the 3D structure modeled from the simulated ensemble Hi-C from model #100. Each dot in the righxt panel represents a 10kb-length particle, and the dots with same color belong to the same predicted TAD ensemble. **(d)** Similarities of predicted single-cell TADs between different thresholds and predictors. **(e)** An example of the simulated data. The upper and lower parts of the heatmap represent the simulated reference and single-cell Hi-C data from model #13, *D*=500. Predicted TADs are shown in sawtooth. AMIs between TADs predicted by deTOKI and IS on the two data sets are 0.873 and 0.660, as, respectively. **(f)** Classification based on deTOKI-predicted TADs of models on chr18:50-55Mb and chr18:10-15Mb, mimicking two single cells. Each dot represents a model, *D*=500. **(g)** Number of misclassifications, using predicted TADs. The color codes for thresholds are identical to **(d)**. *: P<0.05, **: P<0.001, NS: not significant, two-sided Mann-Whitney U test.

The deTOKI outperformed the other tools in the following two respects. First, compared to the other tools, the numbers of TADs predicted by deTOKI and GRiNCH were little affected by data sparsity (Fig.2a and Supplementary Fig.3b). Taking chr10 as an example, the largest absolute log2 fold changes (|log2FC|) in predicted TAD numbers between the down-sampled datasets was 0.26 for GRiNCH and deTOKI, while it was 0.80, 1.38 and 2.40 for IS, SpectralTAD and deDoc, respectively (Fig.2a). Second, on single-cell data, deTOKI predicted TADs more accurately than all other predictors. We took the TADs identified with the full data as the golden standard and quantified the accuracy of predictions by the similarity to this golden standard. Two similarity indexes, i.e., adjusted mutual information (AMI)^50^ and weighted similarity (WS)^42^ were employed. The deTOKI values had higher similarity than all of the other algorithms for both indexes in most chromosomes, i.e., in 19 and 16 out of 22 chromosomes for AMI and WS, respectively (Fig.2b). We also employed two differences indexes, i.e., BP score (BP)^51^ and variation of information (VI)^38^. Although IS performed best with the two indexes (Supplementary Fig.3a), deTOKI performed comparably well in all chromosomes (median BP = 0.49 and 0.51; median VI = 1.58 and 1.61, for IS and deTOKI, respectively).

Moreover, when we performed an additional assessment with binsizes 20kb and 80kb, deTOKI performed equally well as 40kb we described above and better than other tools (Supplementary Fig.3b-c). Finally, marks of the characteristic structural protein CTCF, or histones were found to be enriched in deTOKI-predicted TAD boundaries. Compared to the genomic background, ChIP-seq peaks of H3K4me3, H3K36me3, and CTCF were enriched at the TAD boundary regions predicted by deTOKI, IS and deDoc, while such enrichment was barely seen in those predicted by GRiNCH and SpectralTAD (Supplementary Fig. 3d). These observations further supported the accuracy of the predictions. Taken together, our assessments suggest that deTOKI can stably and accurately predict TADs with ultra-low resolution (i.e., at single-cell level) Hi-C data.

### deTOKI worked well with simulated single-cell Hi-C data

To mimic cell to cell variation, we simulated a single cell Hi-C experiment. The simulated data were generated according to the following protocol. First, we simulated chromosome structures for single cells. By applying a widely used 3D structure modeling tool known as IMP on the bulk Hi-C data, we modeled a 3D chromosome structure ensemble containing about 100 physical chromosome structure models where each model represented a single cell (Fig.2c)^52^. To simplify the simulation, we assumed that each modeled structure in the ensemble would be evenly distributed in the cell population. We have randomly chosen a 5Mb length genome region, i.e., chr18:50-55Mb, to be presented as an example here. To generate single cell Hi-C data from the physical 3D model, we defined the Hi-C contacts as pairs of genome loci with a Euclidean distance less than a threshold *(D)* in the model. Hi-C reads were then sampled from the contacted genomic loci by a binomial distribution. In this work, we tested three threshold *D*s, i.e., 500, 750 and 1000, representing the 20%, 40% and 60% quantiles, respectively, in the distribution of distances between all the genomic loci pairs in the physical model (Supplementary Fig.3g). To define the true TAD domains (reference) in the given single cell, a sufficient number of reads were sampled from the physical model with the sampling probability function of a pair of interacting genome loci being inversely proportional to their Euclidean distance (see Methods). The expected number of reads sequenced from the loci was calculated by normalizing all weights in a genome-wide manner, and the Hi-C reads were then sampled from those genome loci by the Poisson distribution.

The deTOKI can accurately predict TAD structures in simulated single-cell Hi-C data. With the method described above, we simulated the structures of a 5MB region in 100 single cells and generated about 1000 and 0.35M Hi-C contacts each single cell and the reference Hi-C, respectively. By comparing the accuracies (i.e., AMI and WS) of the predictions of the three predictors, we found that when estimated by AMI, deTOKI had significantly higher accuracies than the other tools (Mann-Whitney U test, P < 0.001) for D = 500 and 750. With D=1000, IS had the best performance (Fig.2d); however, the median values of AMI and WS were similar for IS and deTOKI (AMI median = 0.715 and 0.708; WS median = 0.789 and 0.753, respectively). We also employed BP and VI to measure the differences between the predicted TADs and the reference. deTOKI and IS also performed best with these two indexes (Supplementary Fig.3f). This pattern was also seen in an additional randomly selected genome region (chr18:10-15mb, Supplementary Fig.3e, h). For example, in model #13 and with D=500, the deTOKI identified TADs in simulated single cells matched very well with the associated reference Hi-C, while several major TADs were mislabeled using the IS predictor (Fig.2e).

Single cells could also be accurately classified by deTOKI-predicted TADs. As an example, we took the two 5Mb regions of chr11 to represent two type of cells, because being separated by 40Mb on the chromosome they would have few connections. For the simulated 100 models of the two regions (representing two cell types) and using WS as distance, deTOKI-predicted TADs had a better classification power for distinguishing the two cell types than all other tools except SpectralTAD (Fig.2f). Furthermore, the total number of misclassified cells of deTOKI was lower than that of IS, which also performed comparably well in above part (Fig.2g). The success of deTOKI as predictor on the simulated data encouraged us to further assess if the tool would work equally well on experimental single-cell Hi-C data.

### deTOKI predicts TADs with experimental scHi-C data

Next, we compared predictions with three experimental scHi-C datasets, hereafter denoted as Flyamer’s, Tan’s and Li’s dataset (Supplementary Table 1-3)^49,53,54^ We only compared deTOKI with IS and deDoc as the latter two were show to perform relatively well with the simulated sparse data above. We found deTOKI’s predictions to be both more accurate and more stable than those of the two other tools.

First, deTOKI predicted TADs with higher modularity and lower structure entropy. The modularity and structure entropy of a network have previously been used to infer the topological properties of TADs from the Hi-C contact matrix^41,42^. A better defined TAD set is expected to have smaller structure entropy^42^ and larger modularity^41^. With Tan’s and Flyamer’s datasets, deTOKI predicted TADs with lower structure entropies and higher modularities than those of TADs predicted by IS or deDoc (Fig.3a, Supplementary Fig.4a). For example, when we compared the predictions of IS and deDoc in Tan’s data against deTOKI-predicted TADs, we found that TADs in chr1 had the higher modularity and lower structure entropy in all cells, respectively (Fig.3a). In Li’s data, deTOKI also performed best of the three predictors, having the highest modularity and lowest structure entropy in 96 and 73 cells, respectively (Supplementary Fig.4a).

**Fig. 3.**
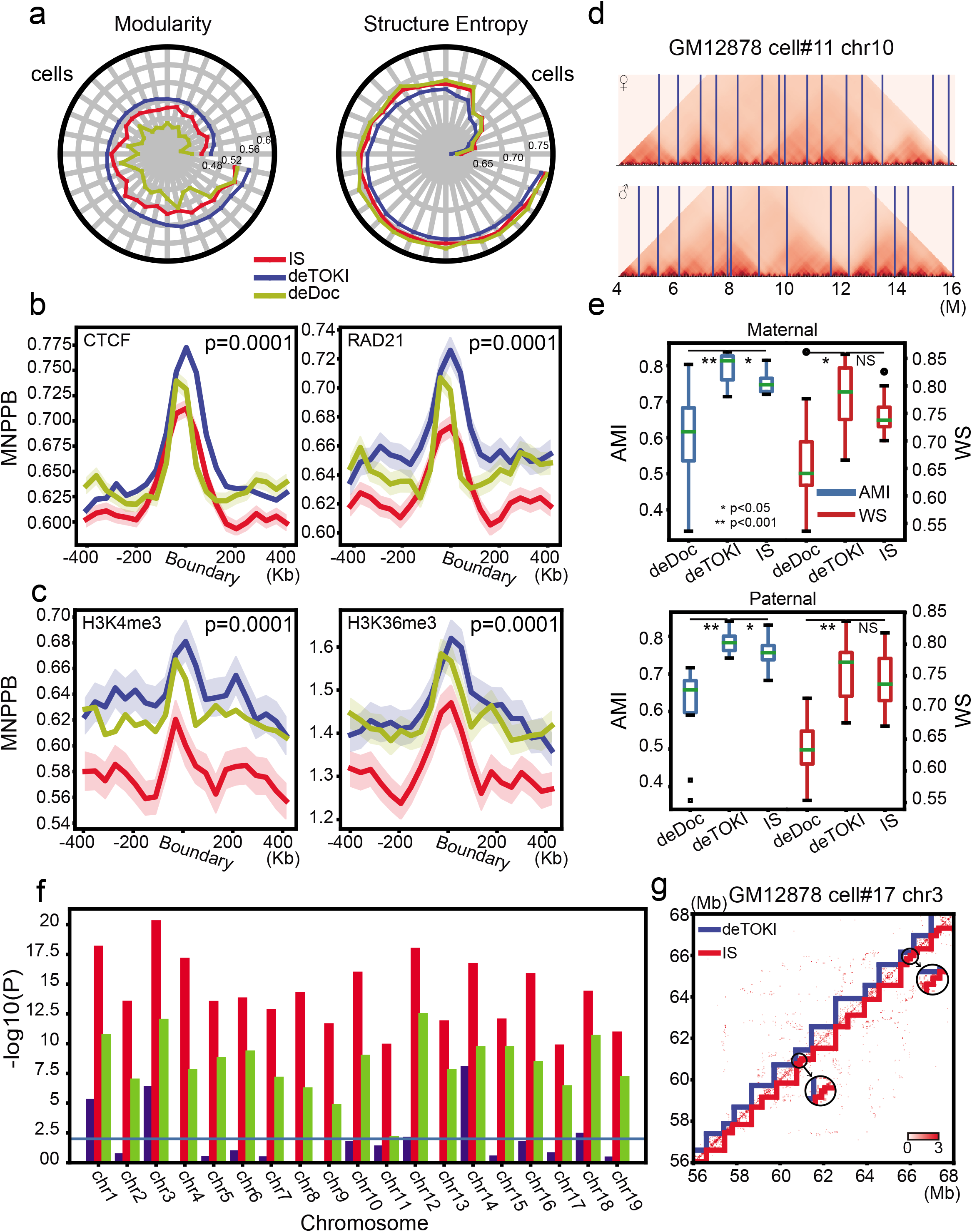
Comparison of TAD callers with Tan’s data^53^ **(a)** The radar plot shows the modularity and structure entropy of predicted TADs in chr1 in 32 cells (15 GM12878 cells and 17 PBMC cells). The cells were ordered by the modularity and structure entropy of deTOKI’s predictions. (b,c) Genome-wide distribution of ChIP-seq peaks of structural proteins (CTCF and RAD21) and histone marks (H3K4me3 and H3K36me3) flanking the single-cell TAD boundaries for the 16 GM12878 cells are shown in **(b)** and **(c),** respectively. The shadows represent 95% confidential intervals as calculated by bootstrap. The *y*-axis represents the mean number of peaks per bin with the same distance to the predicted TAD boundaries (MNPPB). The enrichment p values are calculated by the permutation test (n=10000). **(d)** deTOKI-predicted TAD boundaries match the boundaries predicted by 3D modeling^53^. The example shows the original matrix of radii of gyration for chr10, cell #11 of GM12878. The deTOKI-predicted allelic TAD boundaries are marked with vertical blue lines. **(e)** The AMI and WS between the 3D-modeled hierarchical domains (as defined in^53^) and predicted allelic single-cell TADs in GM12878. **(f)** Significance levels of Pearson’s correlation coefficients between the number of contacts and the number of predicted TADs by deTOKI, deDoc, and IS in each chromosome of 150 mESCs^54^ The threshold “P value = 0.01” is indicated by the horizontal blue line. **(g)** An example of mini-TADs predicted by deTOKI and IS. The mini TADs in the circles are zoomed in as embedded sub-plots. The color codes for the three TAD predictors are all identical to those in **(a)**. *: P<0.05, **: P<0.001, NS: not significant, two-sided Wilcoxon rank-sum test.

Second, the structural protein and histone marks were more enriched at the deTOKI-predicted TAD boundaries. By aggregating the ChIP-seq signals at the predicted TAD boundaries in all single GM12878 cells, we found that the deTOKI-predicted TAD boundaries had higher enrichment of CTCF and Rad21(cohesin) compared to IS and deDoc (Fig.3b). This was also true for H3K36me3 and H3K4me3, the two histone marks previously reported to be enriched in the ensemble TAD boundaries (Fig.3c)^14^.

Third, deTOKI-predicted single-cell TADs were more consistent with the modeled physical structures. Xie and colleagues modeled the physical structure of the haploid chromosomes of single GM 12878 cells at 10kb resolution and proposed an algorithm to infer the chromosome domains from the hierarchical physical structure^53^. Using this haploid physical model and algorithm, we inferred the chromosome domains in a randomly chosen genome region (chr10:4-16M, see Methods). Compared with the deTOKI-predicted haploid single-cell TADs, we found that deTOKI-predicted single-cell TAD boundaries matched the 3D modeling very well (Fig. 3d). Using AMI and WS as the indexes, we compared the 3D-modeled hierarchical domains with the TADs predicted by the three predictors^53^ (Fig.3e). In both maternal and paternal chromosomes, the AMIs of deTOKI’s prediction were significantly higher than those predicted by IS (P = 0.04 and 0.02, respectively, two-sided Wilcoxon rank-sum test). The WS of deTOKI’s prediction was also higher than that predicted by IS (P = 0.1 and 0.35, respectively). As the total number of cells and TADs in this comparison was small, i.e., about 15-25 TADs, we think the significance of the WS was acceptable.

Last, deTOKI exhibited a more stable performance compared to IS. Using chr1 in the PBMC cell #14 as an example, we performed 20 rounds of 50% down-sampling on the single-cell Hi-C reads and predicted TADs from the down-sampled data. Overall, the predictions of both predictors remained largely intact. For example, the distribution of TAD lengths remained similar between the full and the 50% down-sampled data (Supplementary Fig.4b and c). In terms of AMI and WS, deTOKI and IS performed equivalent, i.e., AMI=0.90, WS=0.85 and AMI=0.90, WS=0.87, respectively (Supplementary Fig.4d and e). The AMI and WS of deDoc were 0.80 and 0.71, which is lower than those of deTOKI. However, deTOKI outperformed IS in two respects. First, the number deTOKI-predicted TADs relied much less on reads coverage compared to IS. With Li’s data, the number of IS-predicted TADs was strongly correlated with the reads coverage on all chromosomes, while for deTOKI, only a moderate correlation in this respect was found on five out of nineteen chromosomes (Fig.3f). Second, IS-predicted more questionable mini-TADs, i.e., length < 100kb. The mini-TADs were typically found in the ultra-sparse region, depending on the reads coverage. Within all the TADs, there were 0.29% and 9.81% were considered as mini-TADs, as predicted by deTOKI and IS (Fig.3g and Supplementary Fig.4f and g), in the single-cell, respectively. Taken together, our assessment suggests that deTOKI works fine with experimental single-cell Hi-C data.

### TAD structure is highly dynamic at the single-cell level

Using AMI as the index, we investigated the similarity of TADs between single cells and the ensemble (cell-to-ensemble) and between individual cells (cell-to-cell). By comparing the AMIs of cell-to-ensemble with cell-to-cell in Tan’s data (GM12878 single cells, chr1), we found that TADs in single cells are more similar to the ensemble than TADs between individual cells (Fig.4a, Supplementary Fig.5a-b). In other words, cell-to-ensemble AMIs are significantly higher than cell-to-cell AMIs in all three scHi-C datasets tested. Intriguingly, the average cell-to-cell AMI is even smaller than the cell-to-ensemble AMI of another cell type, e.g., single cells of GM12878 *vs*. ensemble of K562 (Fig.4a). For example, the AMI of cell (GM12878)-to-ensemble (K562) and cell-to-cell (GM12878) AMI are 0.858 and 0.848, respectively (two-sided Wilcoxon rank-sum test, P<0.001, Fig. 4a). Thus, our data suggested that the TAD structure in single cells is quite dynamic, even bigger than inter-cell-type variation. The pattern we showed above is not specific to GM12878, as it can also be seen in the other two tested single-cell Hi-C datasets (Supplementary Fig.5a-b). We note that the average AMI between single cells of GM12878 and ensemble of GM12878 is significantly higher than that of ensemble of K562 (Fig.4a). We tested the assumption that TAD structure carries information for cell identity in the section “Cells can be classified by the TAD structure” below.

**Fig. 4.**
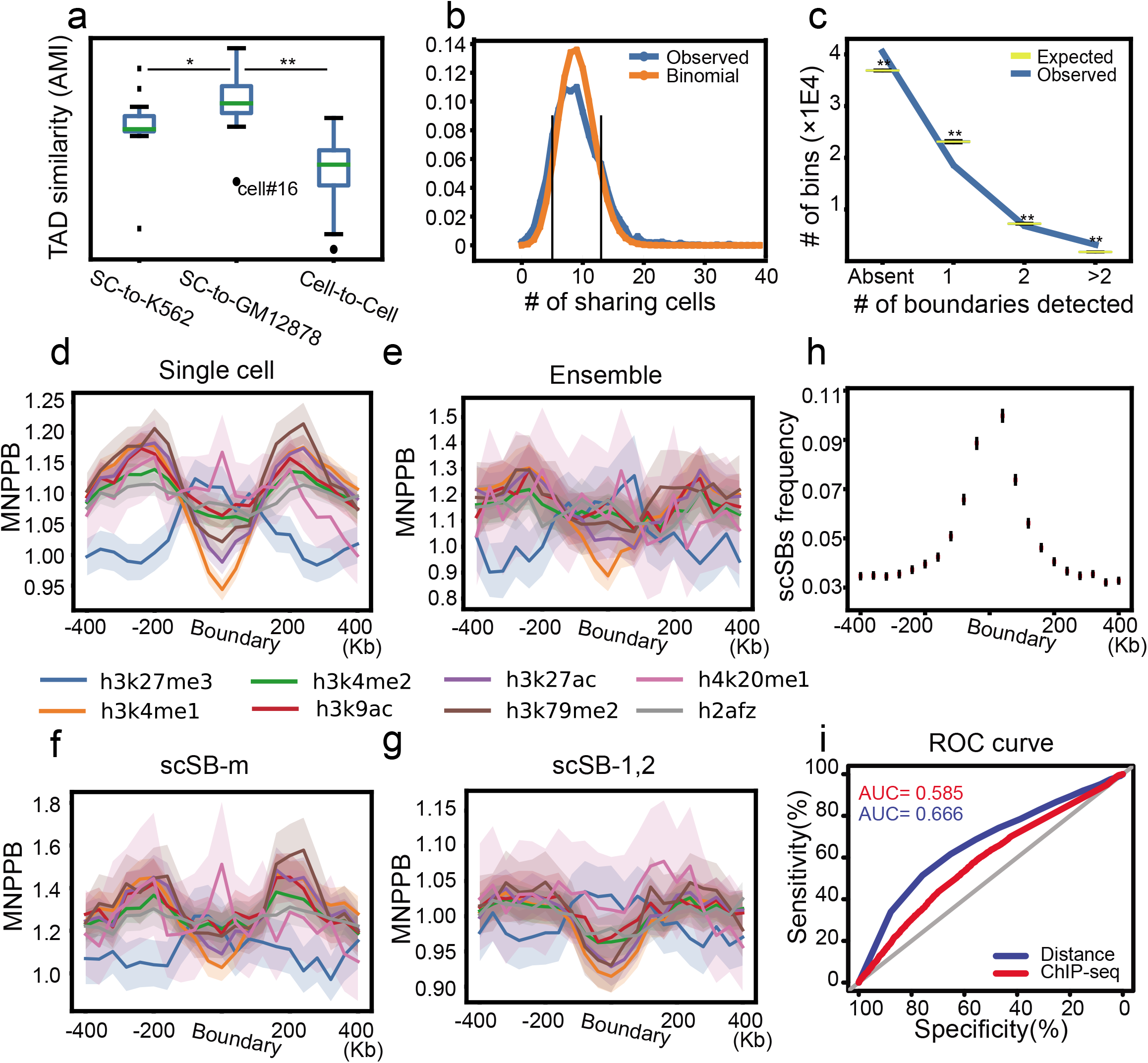
The dynamics of TADs in single cells. **(a)** The cell-to-cell and cell-to-ensemble similarity of the deTOKI-predicted TADs. The single-cell data were from GM12878^53^ and compared to the ensemble in GM12878 and K562. Cell #16 was marked as it was in M/G1 phase. **(b)** The distribution of ensemble TAD boundaries over single cells. The control was set as a binomial distribution under the hypothesis that every ensemble TAD boundary has identical potential of being a single-cell TAD boundary. The vertical black lines marked 13 and 5 indicate the thresholds for the over- and under-represented boundaries in the cell population, respectively. **(c)** The distribution of bins among the classes “scSB-1”, “scSB-2”, “scSB-m”, and “absent”. The permutated control is shown in the boxplots. The distribution of histone marks flanking the deTOKI-predicted single-cell, ensemble, scSB-m, scSB-1 and −2 TAD boundaries are shown in **(d)**, **(e)**, **(f)** and **(g)**, respectively. The *y*-axis of panels **d-g** represents the mean number of peaks per bin with the same distance to the predicted TAD boundaries normalized by the average in the whole genome (MNPPB). The shadows represent 95% confidential intervals as calculated by bootstrap. **(h)** The distribution of scSBs flanking the ensemble TAD boundaries. **(i)** The ROC curves of classification between scSB-1, −2 and scSB-m based on either ChIP-seq peaks or the distance to the nearest ensemble boundaries. *: P<0.05, **: P<0.001, NS: not significant, two-sided Wilcoxon rank-sum test.

Considering that TADs are conserved between cell types^14^, two possible scenarios may explain the above high-level dynamics of the TAD structure in single cells. First, each individual cell employs a subset of the ensemble TADs. Second, each cell has a certain number of additional cell-specific TADs. To test which scenario is the more prevalent in the cell populations tested, we roughly defined three types of variations between TADs, namely, merge, split and shift (see Supplementary Fig. 5c and Methods), where merge does not generate novel TAD boundaries, while split and shift do. Using chr1 as example, we found, on average, 31.8%, 22.3% and 26.6% of merge, split and shift TADs, respectively (Supplementary Fig.5c-e), implying that a notable number of TAD boundaries do not appear in the ensemble TAD structures. We term such boundaries as single-cell-specific boundaries (scSB). In the next two sections, we will sequentially discuss the dynamics of ensemble boundaries and scSBs.

### Unnested ensemble TADs were frequently seen in single GM12878 cells

We asked whether the ensemble TAD boundaries were purely randomly distributed in single cells. A simple assumption for this randomness would be that the distribution of the ensemble TAD boundaries is binomial in the cell population. To examine this assumption, we chose Li’s data as an example and modeled the distribution with a binomial *B*(150,0.06)^54^, where the parameter 0.06 is the average frequency with which an ensemble TAD boundary appears in a single cell. We found that 453 and 452 boundaries (out of 2602) appeared in more than 12 and in less than 6 cells, respectively (Fig.4b). Those numbers significantly deviate from the expectation of binomial null hypothesis (P<0.001). This finding suggests that a group of ensemble TAD boundaries, termed as popular boundaries, occurs more frequently in the cells, while another group of boundaries, termed as unpopular boundaries, tends to be specific to a subpopulation of the cells. GO analysis showed that genes close to the popular boundaries are enriched for terms related to cellular responses to DNA damage stimuli (P=2.21E-3), while genes close to the unpopular boundaries are enriched for terms related to negative regulation of cell-matrix adhesion (P=1.52E-4, Supplementary Fig.6). This result further supported the assumption of a nonrandom distribution of ensemble TAD boundaries in single cells.

Both nested and unnested TAD boundaries were found in the ensemble^26^. We asked how these two types of boundaries are distributed in single cells. We chose chr1 in the GM12878 cells Hi-C data (termed hereinafter as Rao’s data^18^) as an example. We defined the nested and unnested boundaries and compartment domains, as previously described (see Methods^26^). Interestingly, by comparing the number of cells that carry such boundaries, we found that unnested boundaries were significantly enriched in single cells. In the 15 single cells, the 20 nested ensemble TAD boundaries appeared 14 times, while the 20 unnested ensemble boundaries appeared 44 times, being significantly more common than nested ones (P value=0.003, two-sided Wilcoxon rank-sum test, Supplementary Fig.6). Taken together, our analysis suggested that ensemble TADs are dynamic in nature and that unnested ensemble TAD boundaries are more frequently chosen in single GM12878 cells.

### Single-cell-specific TAD boundaries may adhere to the ensemble boundaries

The scSBs may not result entirely from stochastic fluctuation. First, we identified a large number of single-cell-specific boundaries (scSB) using deTOKI. About 89.3% of TAD boundaries in single cells were not found in the ensemble if we defined two boundaries as identical when they were in the same bin. Those scSBs were less likely to result from coverage bias, as strong correlation between the scSBs and read coverage was rarely seen (Cor=0.296, P=0.284). Because of the data sparsity, not all chromosomes found reads in every cell. For this analysis we therefore only looked at the largest chromosome (chr1), for which reads were found in most cells. The following analysis was performed on the whole genome. Second, the distribution of scSBs in the cell population is not random. We grouped all scSBs into 3 classes by the number of cells that carry these scSBs (number=1, =2, >2, denoted as scSB-1, scSB-2, scSB-m, respectively). Compared to the permutated controls, we found that there were far more scSBs in the scSB-m class than in the scSB-1 and scSB-2 classes (P<1E-4, Fig.4c). In fact, most of the genome lacked the potential for TAD boundaries (Fig.4c). A bin is either deficient or relatively prevalent to be a TAD boundary, i.e., either none of the cells, or many cells, took the bin as a boundary, respectively, while very few bins were taken as boundary in only one or two cells.

Third, the scSBs have characteristic histone marks that differ from the ensemble marks. We mapped all histone marks (except for H3K4me3 and H3k36me3) that have publically available ChIP-seq data for GM12878 cells in ENCODE, and we found that the distribution of histone marks showed greater agreement with the single-cell TAD boundaries than with the ensemble TAD boundaries (Fig.4d and e). For example, H3K27me3 and H3K4me1 were found to be enriched and depleted around the boundaries in single cells, while this pattern was much weaker around the ensemble boundaries (Supplementary Fig.7c). We also observed a similar enrichment in IS- and deDoc-identified TAD boundaries (Supplementary Fig. 7a and b). This enrichment of H3K27me3 was higher in the scSB-m than that in scSB-1 and −2 (Fig.4. f and g, Supplementary Fig.7d). Indeed, there were more ChIP-seq peaks representing histone marks in the scSB-m (Supplementary Fig.7e). This line of evidence suggests additional constraint above the stochastic random walk.

To investigate plausible constraints on the scSBs, we compared them with the ensemble boundaries in chr1. We found a strong association between the two classes. First, 7.89% of the scSBs in GM12878 can be found in K562 ensembles, which means that at least some of the GM12878 single-cell-specific boundaries are, likely to be insulative in the K562 ensemble.

Second, the bins that carry scSBs tend to be close to ensemble TAD boundaries. 18.9% of the scSBs are located within an 80kb (+/-40kb) region flanking the ensemble boundaries (Fig. 4h), and the average distance to the nearest ensemble boundaries from scSB-m is significantly smaller than that from both scSB-1 and scSB-2 (Supplementary Fig. 7f). Last, we built a simple logistic regression model to distinguish scSB-1 and −2 from scSB-m using the number of ChIP-seq peaks as features, and we found 5 features, including CTCF, H3K4me1, H3K4me2, H3K9ac and H3K36me3, that were most relevant in this respect (Supplementary Fig. 7g). However, the AUC (0.585) was much lower than the AUC (0.666) of a model that directly used the shortest distance to an ensemble TAD boundary as the feature (Fig. 4i), suggesting that distance is the most important factor restricting the biogenesis of scSB. The importance of distance suggests that genesis of scSBs may not be completely at random but rather tends to fall within certain restricted regions common to all or most human cells, and which is, at least to some extent, represented by the ensemble boundaries.

Altogether, our analysis indicates that a large amount of cell-to-cell variations in the TAD structure, the prevalence of cell-specific TAD boundaries in cells, and a large portion of the single-cell-specific boundaries may not purely result from stochastic fluctuation in the single cells.

### Cells can be classified by the TAD structure

Previously, Tan et al. showed that cell types can be classified using single-cell Hi-C data combined with sequence features of the reads^53^ Now we ask whether the TAD structure alone can be used to classify single cells. Using WS as the similarity index for all three single-cell Hi-C datasets, we found that single cells could be correctly classified by the TAD structure alone. Tan’s dataset^53^ consists of two cell types, GM12878 and PBMC. Both deTOKI and IS can completely distinguish the two cell types using the predicted TAD as feature (AUC=1.0, 1.0 and 0.863, for deTOKI, IS and deDoc, respectively, Fig.5a). Flyamer’s dataset^49^ consists of non-surrounded nucleolus (NSN) and surrounded-nucleolus (SN) oocyte cell types, representing transcriptionally active immature and inactive mature oocytes, respectively^49^. The deTOKI could distinguish these two cell types much better than either IS or deDoc (AUC=0.73, 0.66 and 0.52, for deTOKI, IS and deDoc, respectively, Fig.5b). Flyamer’s dataset also consists of zygote-mats and oocytes. The deTOKI distinguished these better as well (AUC=1.0, 1.0 and 0.89, for deTOKI, IS and deDoc, respectively, Supplementary Fig.8e). In Li’s data^54^, the Methyl-HiC data consists of 150 single cells cultured in two different media: 2i and serum. We found that the TADs predicted by deTOKI could also better distinguish cells with different growing conditions than could IS and deDoc (AUC = 0.691, 0.564 and 0.622, for deTOKI, IS and deDoc, respectively, Fig.5c). The classification of cells in different growth media may not be trivial, as the GO analysis showed that the genes close to serum-specific TAD boundaries were enriched for the term “DNA methylation on cytosine” (Supplementary Fig.8a and b, P=7.28E-4), which is consistent with the fact that the serum-cultured mESC has a higher DNA methylation rate^54^ The serum-specific TAD boundaries were also enriched for the gene regulation-related GO terms, e.g., positive regulation of gene expression and epigenetic (P = 8.20E-4), which agrees with the fact that serum-cultured mESCs have more heterogeneous transcriptional activity than cells cultured in 2i^55–57^ Further, epigenetic features were distinguished between the serum- and 2i-specific TAD boundaries (Supplementary Fig.8c). Altogether, deTOKI predicted TADs in single cells carrying reliable information about cell identity.

**Fig. 5.**
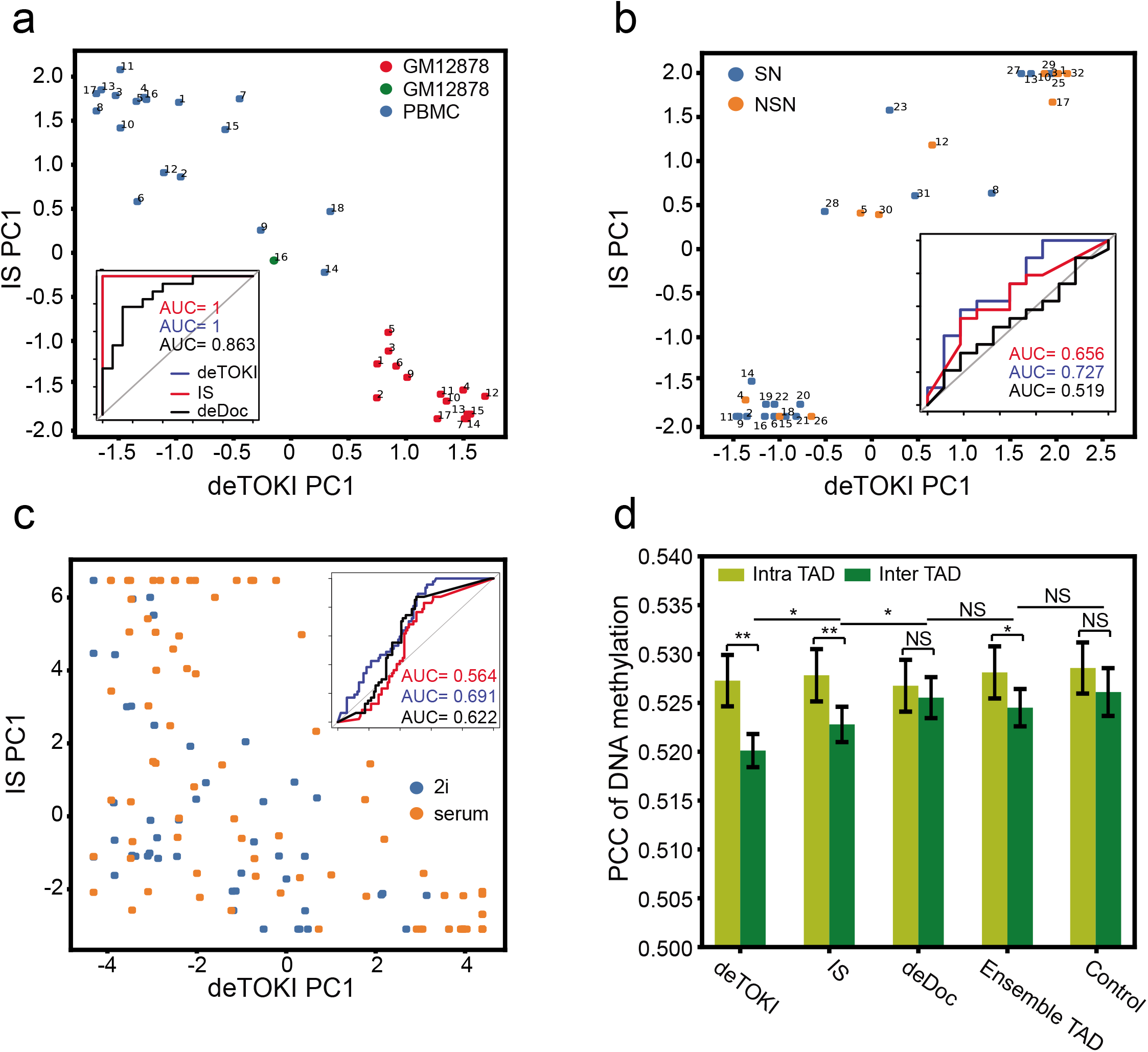
The deTOKI-predicted single-cell TADs characterize cell identity. The classification of single cells based on predicted TAD boundaries in Tan’s, Li’s and Flyamer’s datasets are shown in **(a)**, **(b)** and **(c)**, respectively. Each dot represents a cell. The *x-* and *y*-axis represent the PC1 calculated by deTOKI and IS, respectively. The embedded plots show the AUC of the classification by each program. Cell #16 is colored in green in panel **(a),** as it was in M/G1 phase. **(d)** The correlation of DNA methylation between bin pairs. The “inter TAD” indicates that the two bins are separated by a predicted TAD boundary, while the “intra TAD” indicates that the two bins belong to the same predicted TAD. Only bins fulfilling the criteria of 1) being members of pairs separated by 240Kb *and* 2) having reads containing a total of 2 or more CpGs, were included in this plot. *: P<0.05, **: P<0.001, NS: not significant, Fisher’s z-test^63^

### The DNA methylation pattern is highly correlated between TAD boundaries at the single-cell level

It has been suggested that spatial approximated genome loci are prone to share similar epigenetic patterns^58^ We thus asked if this feature could also be seen in single cells. If this be the case, we would expect to see lower correlations in DNA methylation between the inter-TAD bins than that between intra-TAD bins in the single cells. To test this speculation, we looked at Li’s data^54^. First, at the ensemble TAD level, the genome loci flanking the strongly insulated TAD boundaries have lower correlations on DNA methylation than those flanking the weakly insulated TAD boundaries (Supplementary Fig.8d). We classified the ensemble TAD boundaries into strong and weak groups by their insulation scores and calculated the inter-TAD PCCs for all the boundaries. The average PCCs were 0.546 and 0.490 for weak and strong boundaries, respectively. Next, we examined the inter-TAD and intra-TAD PCCs of DNA methylation level in the single cells. Indeed, when the TADs were defined by deTOKI or IS (Fig.5d), the intra-TAD PCCs were significantly larger than the inter-TAD PCCs, while when the TADs were defined by deDoc or the shuffled control, little difference was noted. The PCCs of inter-TAD bins from deTOKI-predicted TADs were significantly lower than those from IS-predicted TADs (PCC=0.520 *vs*. 0.523, for deTOKI and IS, respectively, p=0.003), implying that the boundaries predicted by deTOKI are more spatially insulated in single cells. Although the average PCC between inter-TAD bins was relatively low, it remains notable. We speculate that this might be caused by the existence of weak TAD boundaries, as discussed above. Together, our analysis suggested that spatially approximate chromatin loci are prone to carry similar epigenetic features and that the dynamic nature of TAD structures at the single-cell level has notable consequences for the ensemble of the epigenetic landscape.

## Discussion

In present work, we have developed a TAD identification algorithm that can work on sparse data at the single-cell level. We assessed the accuracy and robustness of deTOKI in down-sampled, simulated and experimental single-cell Hi-C data, and we compared deTOKI to the two best-performing tools on sparse data, IS and deDoc^43^. The assessment showed that deTOKI not only outperformed IS and deDoc, but also reliably predicted TADs in experimental single-cell Hi-C data and is thus the first published tool with such a capacity.

We took advantage of NMF on handling sparse data for decomposition of the Hi-C contact matrix. NMF has been widely used in single-cell data analysis, e.g., coupled NMF^45^ The boundaries defined by deTOKI were the optimal saddle points, which are also the genome loci that insulate chromatin interactions. The combination of NMF and insulation detection enabled deTOKI to achieve reliable TAD prediction on sparse data.

Future deTOKI work will involve the following features. First, we will improve sensitivity in the contact desert regions. New experimental technologies for higher data coverage have been able to reach the contact desert region^53^, but algorithms can still be improved. Deep neural networks have proven to be a powerful method for pattern recognition. With an increasing growth of single-cell Hi-C data, the deep leaning-based methods are expected to improve the sensitivity in domain calling in the contact desert regions. Second, introduction of a better assessment for TAD reliability would be extremely useful when the detection probes the deep contact desert regions. Finally, deTOKI needs to gain some speed. Although the current running speed of deTOKI is acceptable, it is slower than IS and deDoc. Parallelization is one way to improve the speed, as deTOKI works on split genome fragments. However, we sought to optimize the algorithm so that access to a supercomputer is not necessary to scan the whole genome.

With the ability of probing TAD structures in single cells, we examined the dynamics of the TAD boundaries. Three novel features were revealed. First, although the cell-to-cell variation is large, most single-cell TAD boundaries adhered the ensemble consensus. Since only a small fraction of boundaries in the ensemble can be detected in each single cell, the dynamics of TADs is likely to be high. However, since most scSBs adhered to the ensemble consensus this may indicate the existence of sub-populations in the isogenic cell population. Whether the cells would constantly stay in one sub-population or switch between sub-populations will be an interesting question to ask in future studies. Second, our data showed that TAD boundaries are prone to be unnested TAD boundaries, while little bias was noticed in compartment domain. This result may indicate that the biogenesis of TADs and the compartment domain is different in principle. The last novel feature is the enrichment of certain histone marks at, or flanking, the scSBs, but weaker at the ensemble boundaries (Figure 4d-g). As we do not have single-cell ChIP-seq data available at present, it would be extremely interesting to ask if those histone marks do, indeed, can be observed in single cells. If the answer is in the affirmative, then many as yet undiscovered properties of scSBs may be linked to the function of TADs in single cells. Preliminary GO analysis showed strong association between the enriched functional terms and cell identity (Supplementary Fig. 8), hinting at the profound functions the scSBs may have. To further reveal the mechanisms of 3D genome folding, the principle and function of the domain structure at single-cell level will be key questions to ask. The deTOKI provides a basic tool for addressing those questions.

## Methods and Materials

### The simulation of single-cell and reference Hi-C

To simulate Hi-C, we constructed a 3D model using IMP with default settings at 10kb resolution for any given 5Mb genome region^52^. We simulated the reference and single-cell Hi-C data as follows^59^.

Simulation of the reference Hi-C: For any two genome loci *i* and *j,* the weight was set as

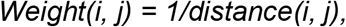

where the distance was Euclidean. The chance of being sequenced in a Hi-C was then set as the normalized weights, and the expected read number was calculated by the chance times the total number of reads. The normalization was performed so that the total number of sequence reads was identical to that of widely used bulk Hi-C data^14^, being the equivalent of 0.35M reads per 5-Mb region. Hi-C reads were simulated by Poisson distribution with the expectation calculated above.

Simulation of the single-cell Hi-C. For any two genome loci *i* and *j,* the weight was set as

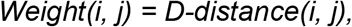

where the distance was Euclidean, and D is the threshold. Only genome loci having an Euclidean distance less than D were considered contacting. The chance of being sequenced in a single-cell Hi-C was then set as the normalized weights, and the expected read number was calculated by the chance times the total number of reads. The normalization was performed so that the total number of sequenced reads was identical to Tan’s data^53^, being equivalent to 1000 reads per 5-Mb region. Hi-C reads were simulated by Binomial distribution with the expectation just calculated. The 40kb resolution Hi-C contact matrix was used for actual TAD detection. Thus, the simulated 10kb resolution matrix was binned into 40kb resolution.

### Processing of Hi-C data

Bulk Hi-C data were normalized by the ICE method^60^, while we did not normalize single-cell Hi-C data owing to its sparse nature, and we also skipped the normalization step on the down-sampled data when its sparsity was comparable to that of single cells. In this work, we used a sampling rate of 1/800 at the single-cell level. All simulated and experimental data used in this study are summarized in supplementary table 4.

### deTOKI

We split the chromosomes into 8-Mb sliding windows overlapping each other by 4Mb (Fig. 1b), and removed windows with less than 100 intra-window contacts. The TADs were then predicted as follows.

1. The clustering of bins. In each 8-Mb window, we perform NMF on its contact sub-matrix by the function “sklearn.decomposition.NMF” of the scikit-learn package in Python^44^, with “random” being the initialization setting (Fig. 1c). The parameter “n_components” represents the dimension of the factor matrices. The parameter “n_components” traverse an appropriate interval according to the average length of the TAD(s) and the length of the window. The suggested numbers for mammalian cells were 8, 9, …, 13. The bin *i* and bin *j* are clustered if the maximums in columns *i* and *j* of the coefficient matrix is the same as in the raw.
2. TAD boundary detections. For each candidate of “n_componets” equal to n, we perform NMF *k* times in which the seed for random initialization, namely “random_state”, traverses within the interval [0, k-1]. The default *k* was 10, as we found little difference on the predicted TADs between *k* = 10 and *50* for both single-cell and bulk Hi-C data (Supplementary Fig.1a). We define a consensus map ***C***, 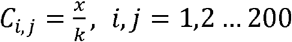, where *x* denotes the number of NMFs that have bin *i* and bin *j* clustered together. Then, the cluster rate of any given bin *i* (*CR_i_*) was defined as the average value of all elements in the sub-square-matrix of ***C*** cornered at *(i, i)* with 11 bins along the matrix diagonal. The location and strength of the bins that have local minimum cluster rates were recorded. The strength of *bin_i_* was defined as the local maximum *CR* minus *CR_i_*. Thus, the TAD boundaries were defined as the strongest *n-1* bins and the points that have a *CR* strength larger than 0.3 (Fig. 1d).
3. The silhouette coefficient calculation. The silhouette coefficient was introduced to provide an evaluation of clustering validity, and it is often used to select an ‘appropriate’ number of clusters^61^. For each candidate of “n_componets”, we calculated the silhouette coefficient between the consensus map *C_i,j_* and the TAD boundaries {*i*_1_,… *i_m_*}, as in the following,

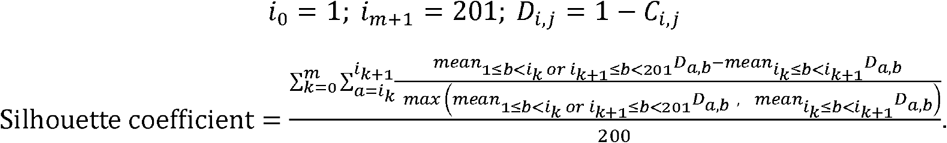 The candidate *n* with the biggest silhouette coefficient was chosen, and its associated domain boundaries were considered as the final prediction (Fig. 1d and e).
4. Suggested parameter settings. For low resolution Hi-C data, we recommend 8-Mb window and 40kb bin-size as the proper setting. Although the difference of predicted TAD between different bin-size for both single-cell and bulk Hi-C data were found minor, they are adjustable as parameters in the deTOKI (Supplementary Fig.1b).
5. Running time. The expected running time is O(N), in which N is the length of contact matrix (Supplementary Fig.1c). It took about one hour to identify TADs in 40 kb resolution data of the whole genome of mm9 with Flyamer’s data^49^ The testing was performed in a computer with Intel(R) Xeon(R) CPU E5-2640 v3 @ 2.60GHz with one core, and it could be as fast as finishing the same job in 6 minutes when using 16 cores (Supplementary Fig.1c).

### Execution of other TAD predictors

Most of TAD predictors were executed with default parameters. We removed the mini-TADs predicted by deDoc, i.e., TADs shorter than 200kb and 300kb, in the simulated and experimental single cell data, respectively. We calculated hierarchical domain and radii of gyration in the single-cell Hi-C data according to Tan et al^53^ To properly compare hierarchical domains and TADs, we cut the hierarchical tree such that the number of domains and TADs were similar.

### The similarity of two sets of TADs

Given two sets of TADs, T = {T_1_, T_2_, …,*T_n_*} and K = {K_1_, *K*_2_, *K^m^*}, we assess their similarity using adjusted mutual information (AMI)^50^, weighted similarity (WS)^42^, BP distance (BP)^51^ and Variation of Information (VI)^38^.

### Adjusted Mutual Information AMI(T, K)

Mutual information MI (T, K) was defined as

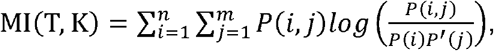

where

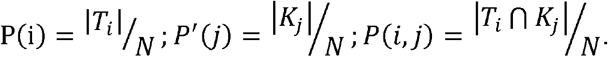

Then, the adjusted mutual information AMI (T, K) was defined as

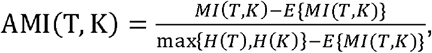

where *H* denotes the standard Shannon entropy, and *E* denotes expectation. AMI was calculated by the function adjusted_mutual_info_score in the Python module sklearn.metrics. In real calculation, all predicted TADs and intermediate windows of TADs are included in *T* and *K*.

### Weight Similarity WS(T, K)

The weight similarity WS(T,K) was defined as

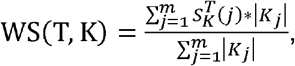

where

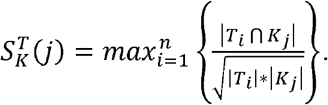

Because WS is an asymmetric index for similarity, we always put the predicted TADs from raw data in *T* and the TADs from down-sampled data in *K*, while the intermediate windows of the TADs were not included in either *T* or *K*.

### The enrichment of ChIP-seq peaks at the boundary region of TADs

For any given resolution, e.g., 40kb, a boundary region was represented by a vector of 21 entries, where the 1^st^ to the 10^th^ entries represent upstream 10 bins, the 12^th^ to the 21^st^ entries represent downstream 10 bins, and the 11^th^ bin represents the middle point of the boundary. The value of each entry is the number of ChIP-seq peaks in each bin, and the middle entry is the total number of peaks in this boundary. These vectors are then summed (entry-wise) up to a total vector *v_i_*(*i* =-10,-9, …,9,10). We define the MNPPB (mean number of peaks per bin) to reflect the enrichment of ChIP-seq peaks on the boundary of TADs, as

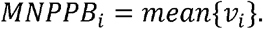

### Evaluation index of TADs

Given TADs *T = {T1, T2, T3, T4,.,.,Tn}* and contact matrix, *F_i,j_*(*i,j* = 1,2 …*N*), we assess the structure property of TADs by using the following two indices, according to the literature.

### Structure Entropy (SE)^42^

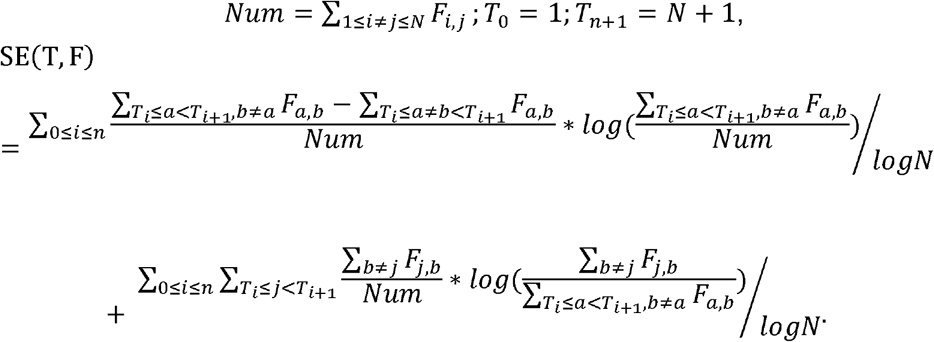

### Modularity Index (M)^41^

After removing the entries in the diagonal of the contact matrix, we split the chromosome into 6-Mb nonoverlapping windows. We further removed windows with less than 100 intra-window contacts, as the method was designed for TAD assessment with sufficient data^41^. For each window, we consider TAD boundaries in/of the region *S = {S0=0, S1, S2, S3, S4,,.,Sm=150}* and the log transform contact matrix of the region *E_i,j_*(*i,j* = 1,2 …150). Then we calculated the modularity of this region as follows. The modularity of each 6-Mb region was then averaged into a modularity index.

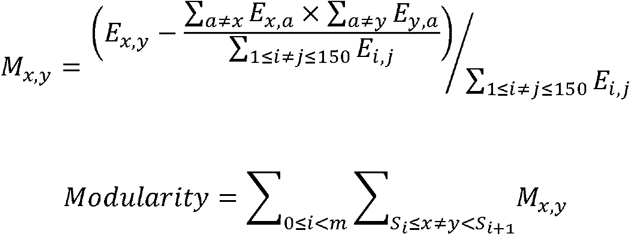

### Unsupervised classification of single cells

The classification based on TAD similarity in chromosome k between n cells is performed by PCA of the *C_i,j,k_*(*i,j* = 1,2, …,*n*), which is the self-Spearman correlation coefficient matrix of the similarity matrix *M_i,j,k_* (*i,j* = 1,2, …,*n*), calculated as follows:

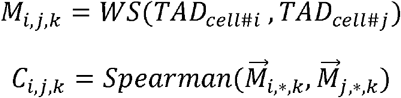

The classification based on the TAD similarity in all chromosomes between cells is performed by PCA of the matrix *T_i,j_*(*i,j* = l,2,…,*n*), calculated as follows:

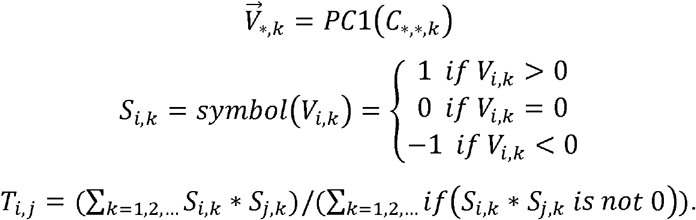

If data in chromosome *k* of cell *i* are not available, then *S_i,k_* will be set to 0, and the *T*s can only be calculated when both cells have sufficent data available. After we have calculated PC1 of matrix *T_i,j_*(*i,j* = 1,2, …,*n*), we get the classification index of each cell. Last, we access the classification by calculating the AUC of the ROC curve.

### The definition of matched, merged, split and shifted TADs (Supplementary Figure 5c)

As previously defined^62^, the TAD boundary regions were defined as the flanking 100kb region of the boundary bins, and the region between these TAD boundary regions was considered as being inside the TADs. Matched TADs: if both boundaries of a TAD in one condition aligned within TAD boundary regions in another condition. Merged TADs: if two or more TADs in one condition aligned inside of a TAD in another condition. Split TADs: if one boundary aligns to a boundary region of one TAD and the other boundary aligns inside of a different TAD. Shifted TADs: if the two boundaries of a TAD align into two different TADs.

### The definition of nested TAD boundary, unnested TAD boundary, and compartment domain boundary

Using Rao’s data at 40kb resolution in chromosome one^18^, we called TADs with deTOKI and obtained 253 TADs. Then the TAD boundaries were sorted by the number of contacts between the up- and downstream 400kb regions. The top 20 TAD boundaries with highest cross-boundary contacts were defined as nested TAD boundaries, and the 21^st^ to the 40^th^ were defined as unnested TAD boundaries. A compartment domain boundary was defined as the TAD boundaries that had different compartment scores between the flanking bins.

### Definition of 2i-specific TAD boundary and serum-specific TAD boundary

For 103 serum cells and 47 2i cells^54^, we defined the bias of one bin as the difference in proportion between the serum cells and the 2i cells for which this bin represents a TAD boundary. We sorted all 40kb bins in whole genome according to the bias value and defined the top and bottom 400 bins as serum- and 2i- specific TAD boundaries, respectively.

## Availability

The source code can be freely accessed through github at https://github.com/lixiaoms/TOKI.

## Funding

This work was supported by Beijing Natural Science Foundation (Z200021), Special investigation on science and technology basic resources of the MOST, China (2019FY100102), the Beijing Advanced Discipline Fund (115200S001), the Strategic Priority Research Program of the Chinese Academy of Sciences, China (XDA24020307), the National Key R&D Program of China (2018YFC2000400), and the National Nature Science Foundation of China (31671342, 31871331, 91940304).

## Authors’ contributions

XL and ZZ conceived this project. XL performed the experiments, analyzed data, XL and ZZ prepared the manuscript. All authors read and approved the final manuscript.

## Competing interests

The authors declare that they have no competing interests.

## Acknowledgements

We thank Dr. Bingxiang Xu for his help in discussions at the early stage of this project. Dr. Geir Skogerboe for helpful discussion and language proofreading, Mr. David Martin performed English language editorial services.

## Notes

### Competing Interest Statement

The authors have declared no competing interest.

