## Supplementary figures and images for "DeTOKI identifies and characterizes the dynamics of chromatin topologically associating domains in a single cell"

### fig s1

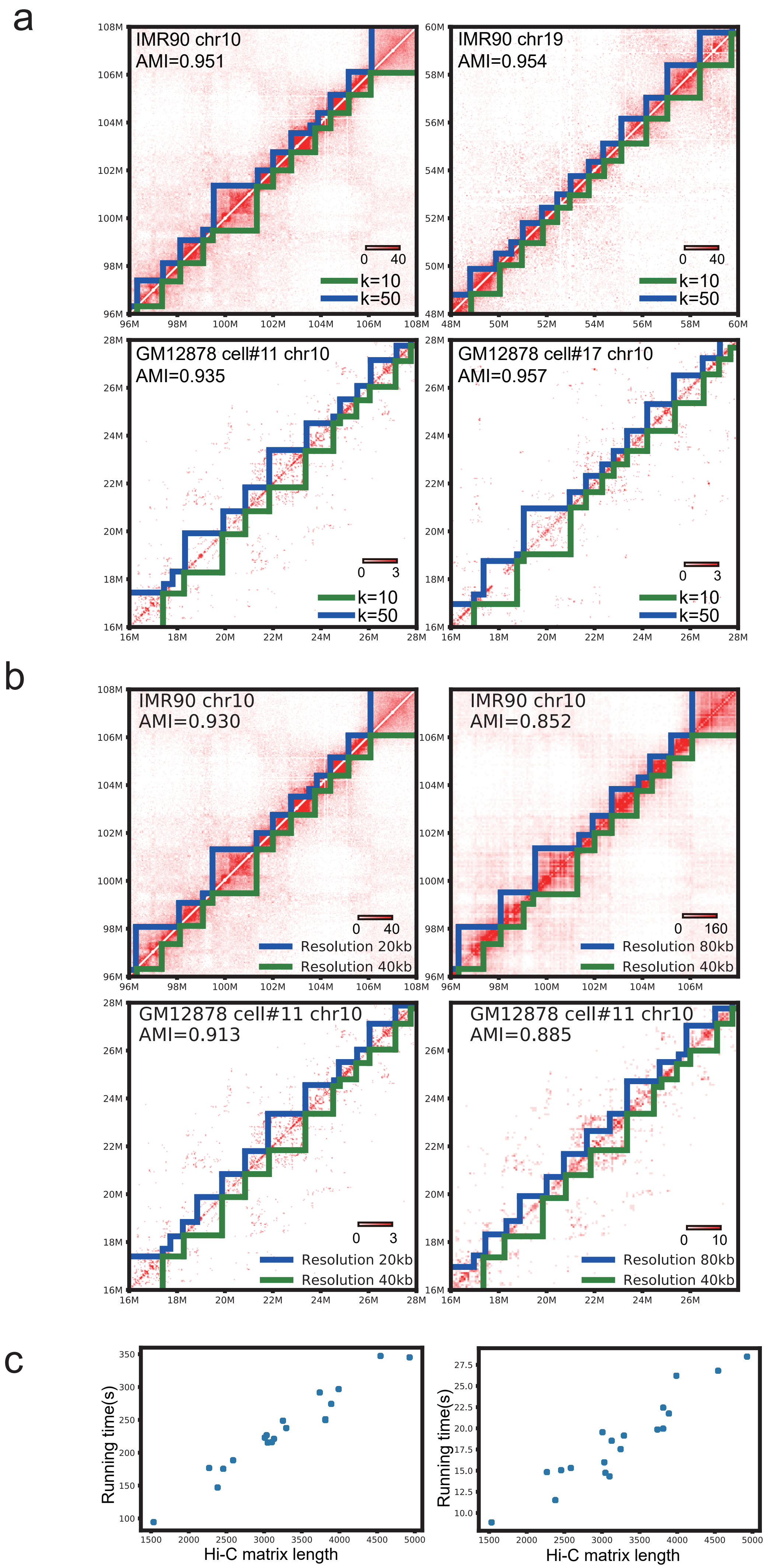

### fig s2

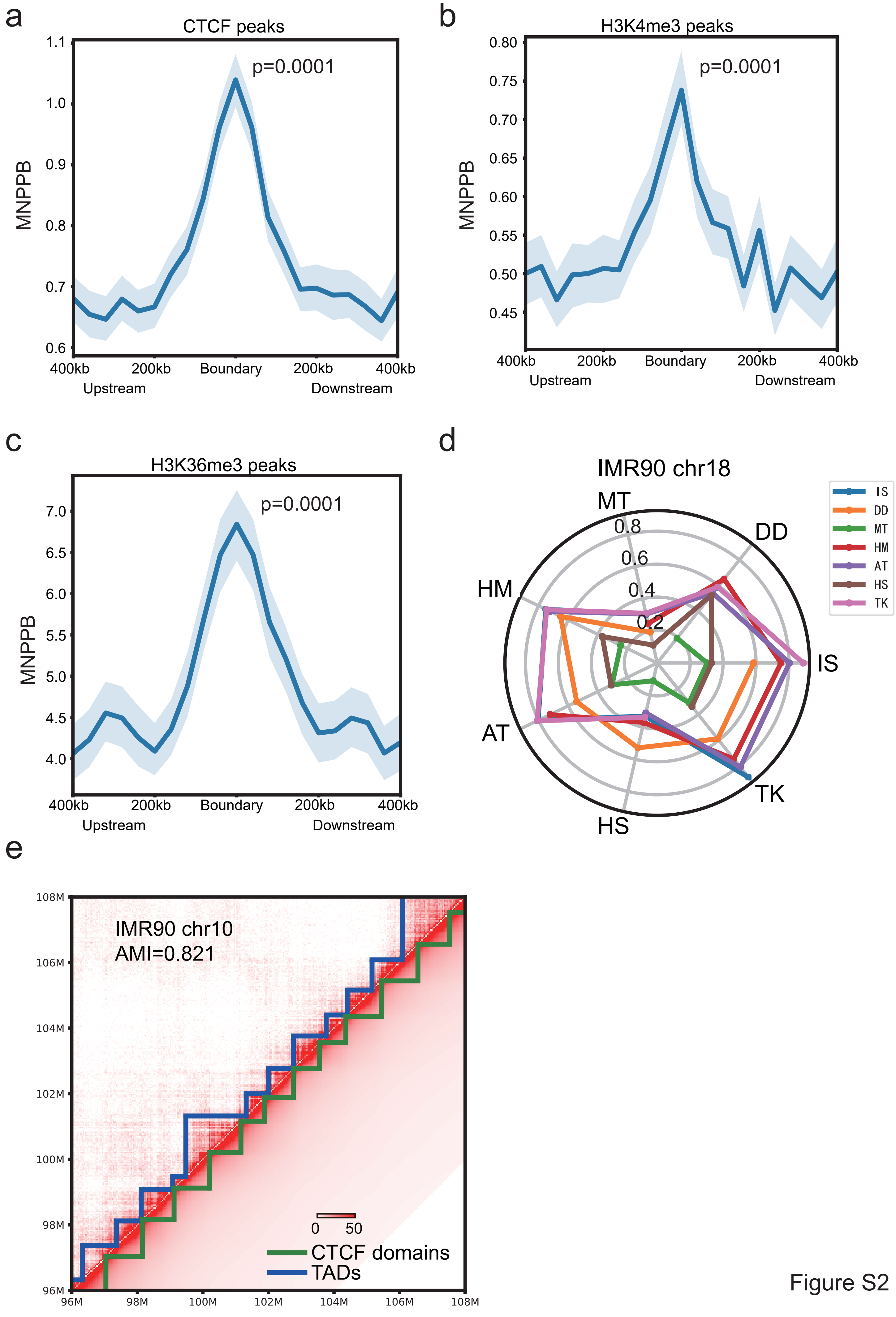

### fig s3

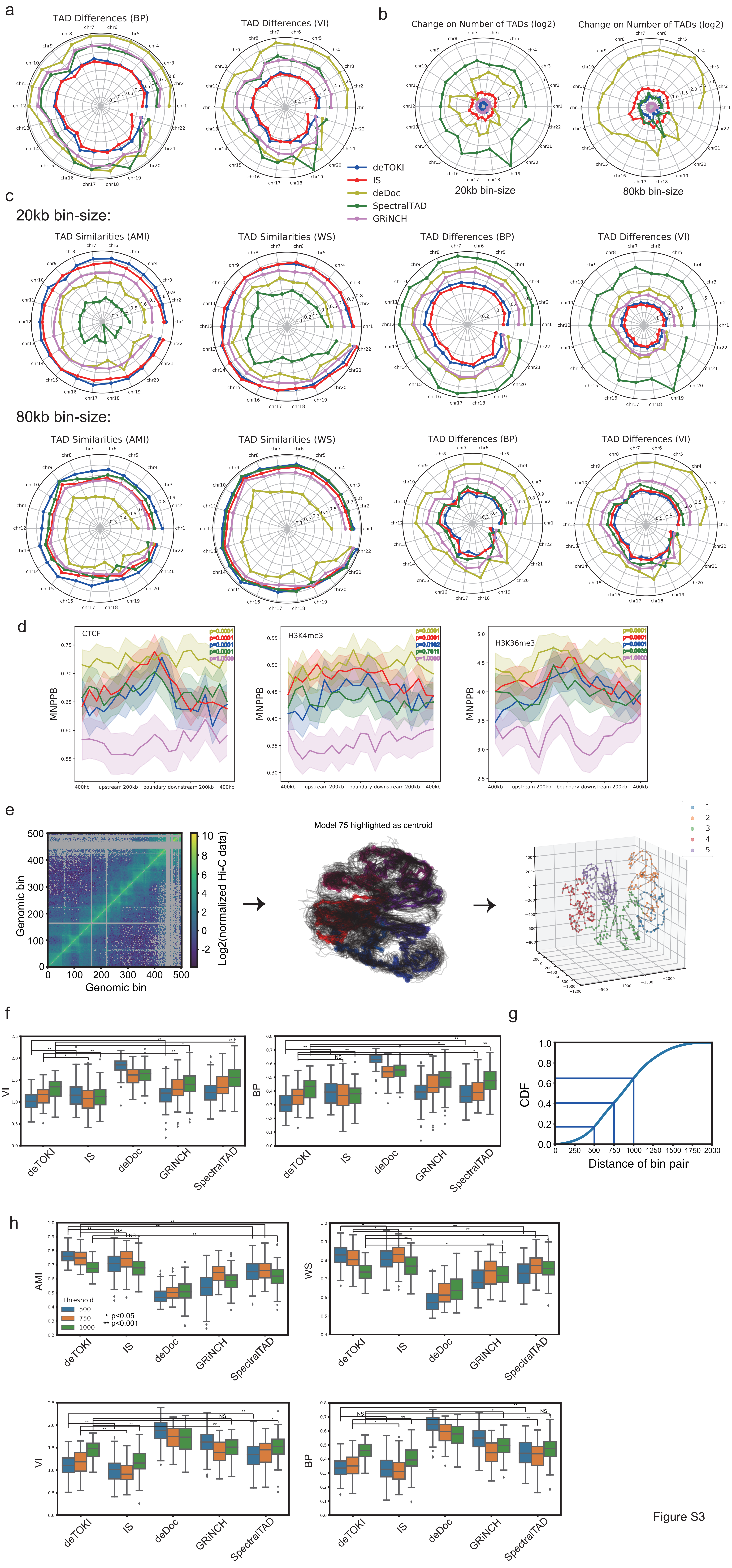

### fig s4

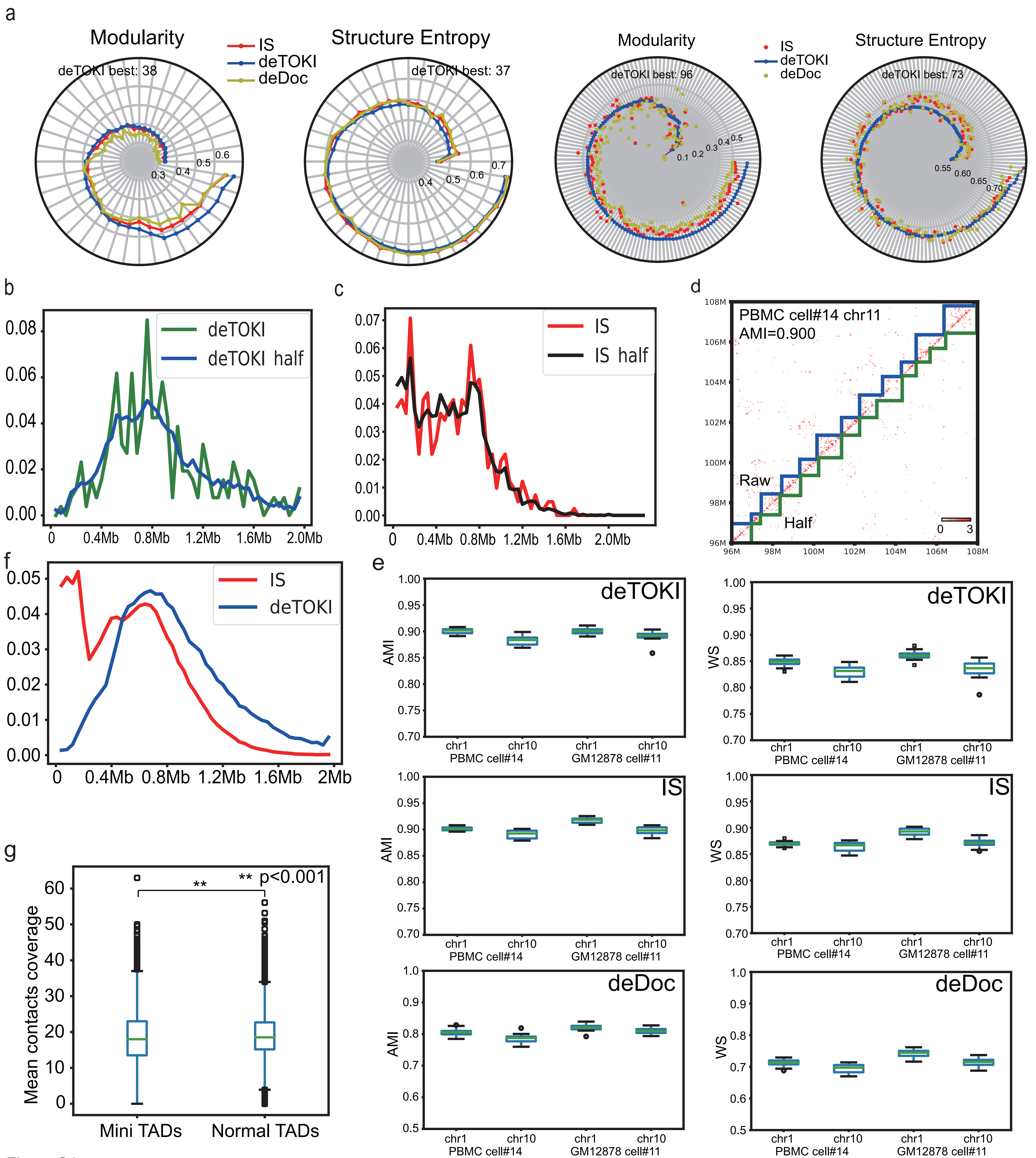

### fig s5

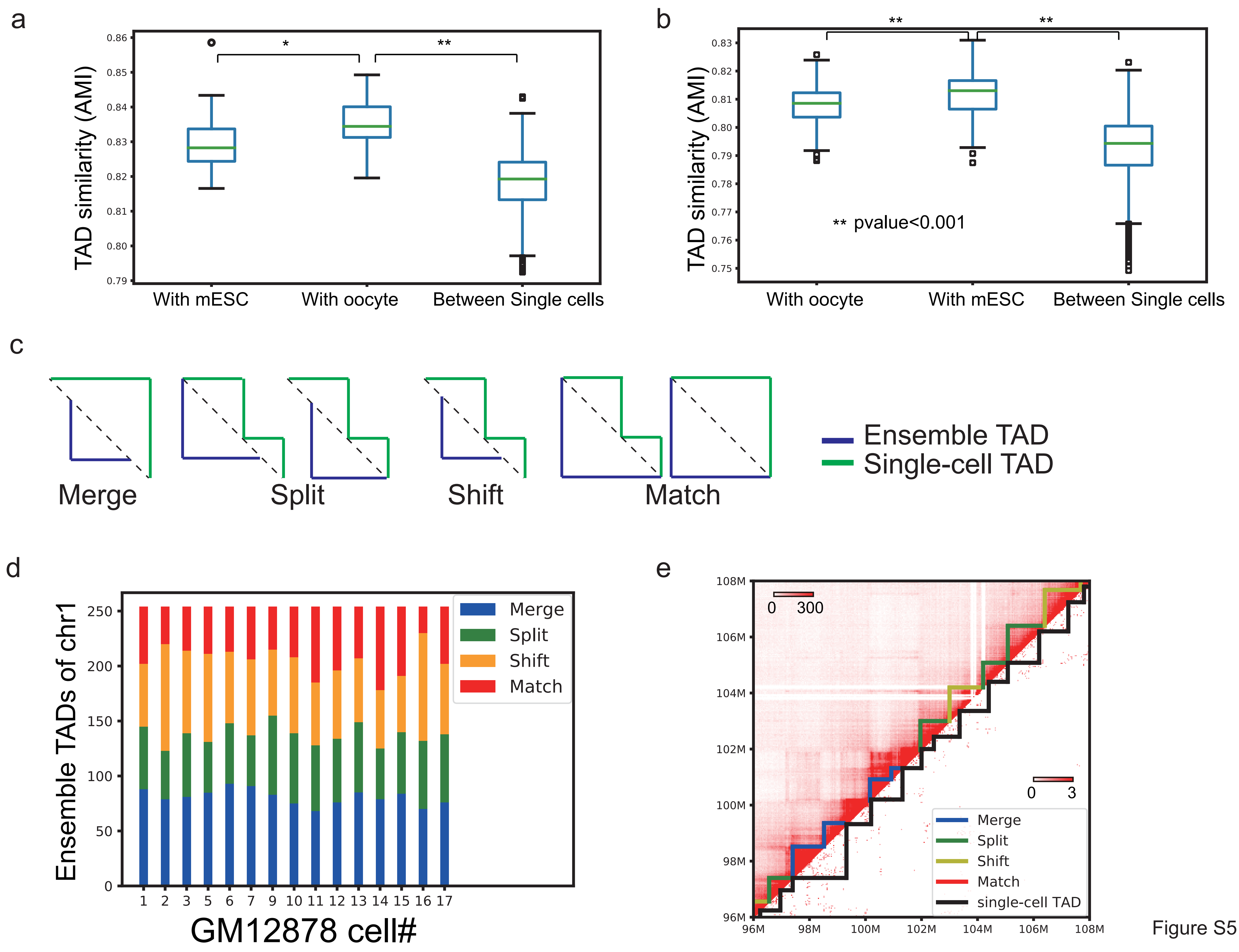

### fig s6

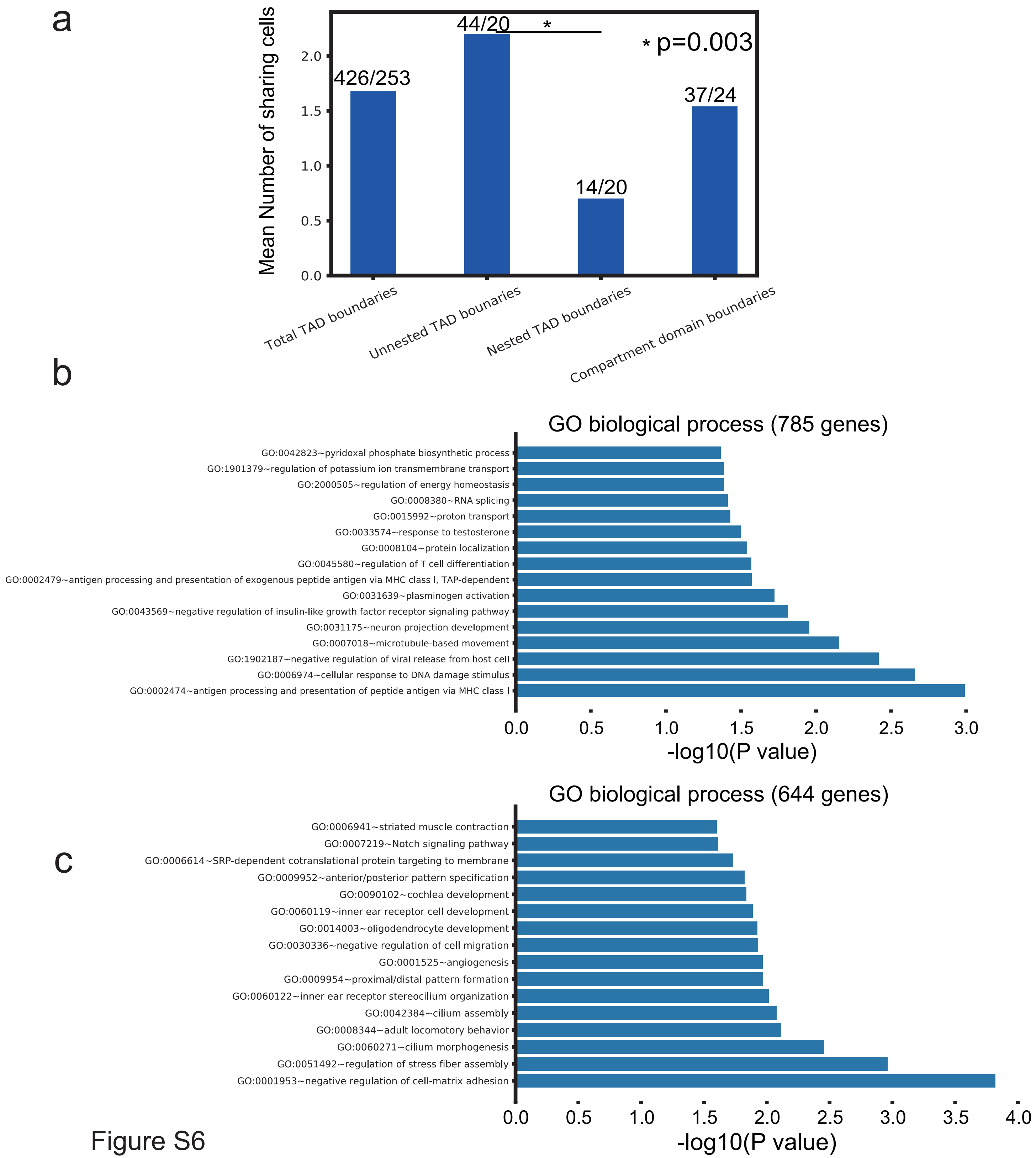

### fig s7

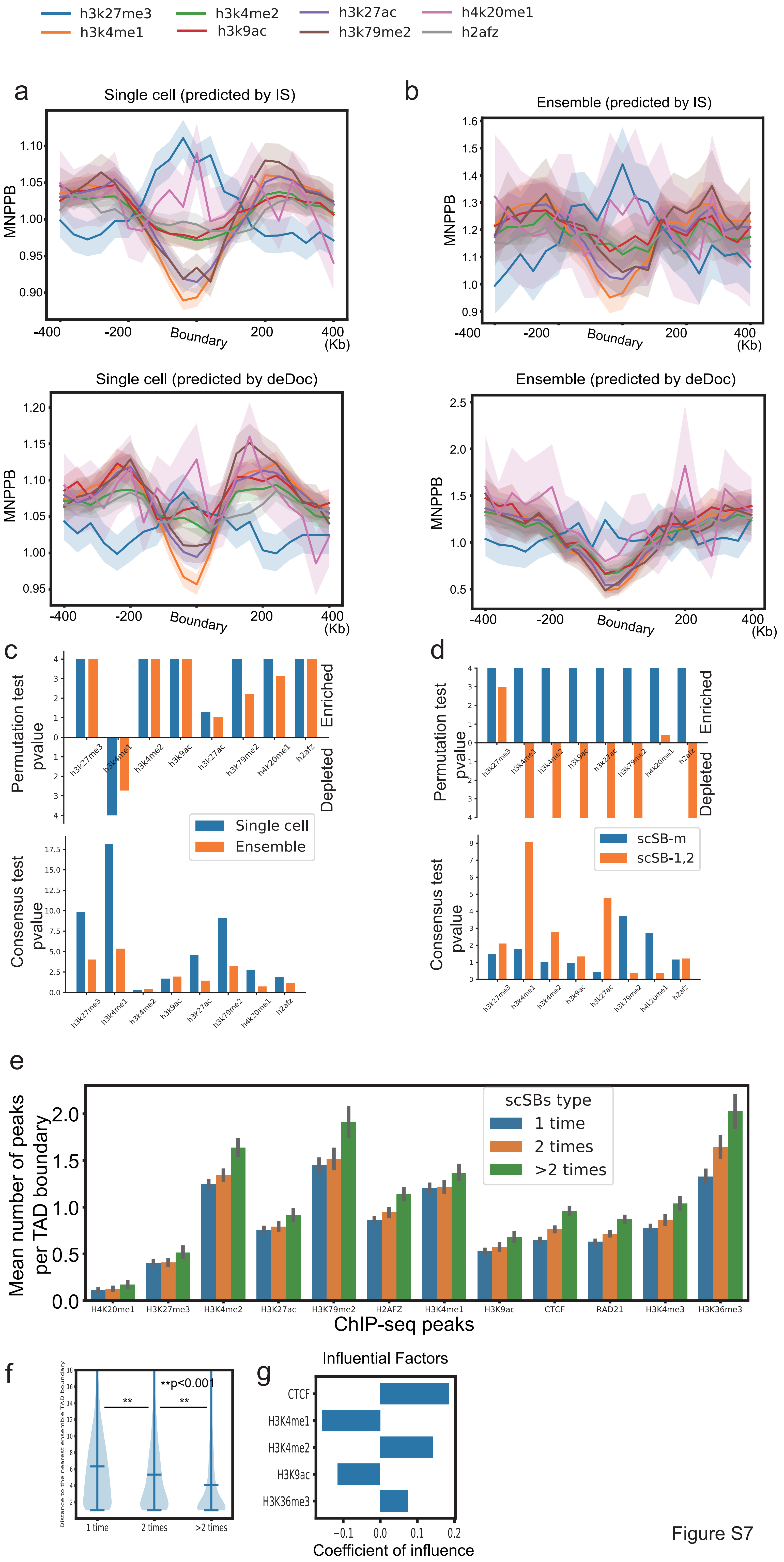
