## Supplementary material for "DeTOKI identifies and characterizes the dynamics of chromatin topologically associating domains in a single cell": fig s8

a

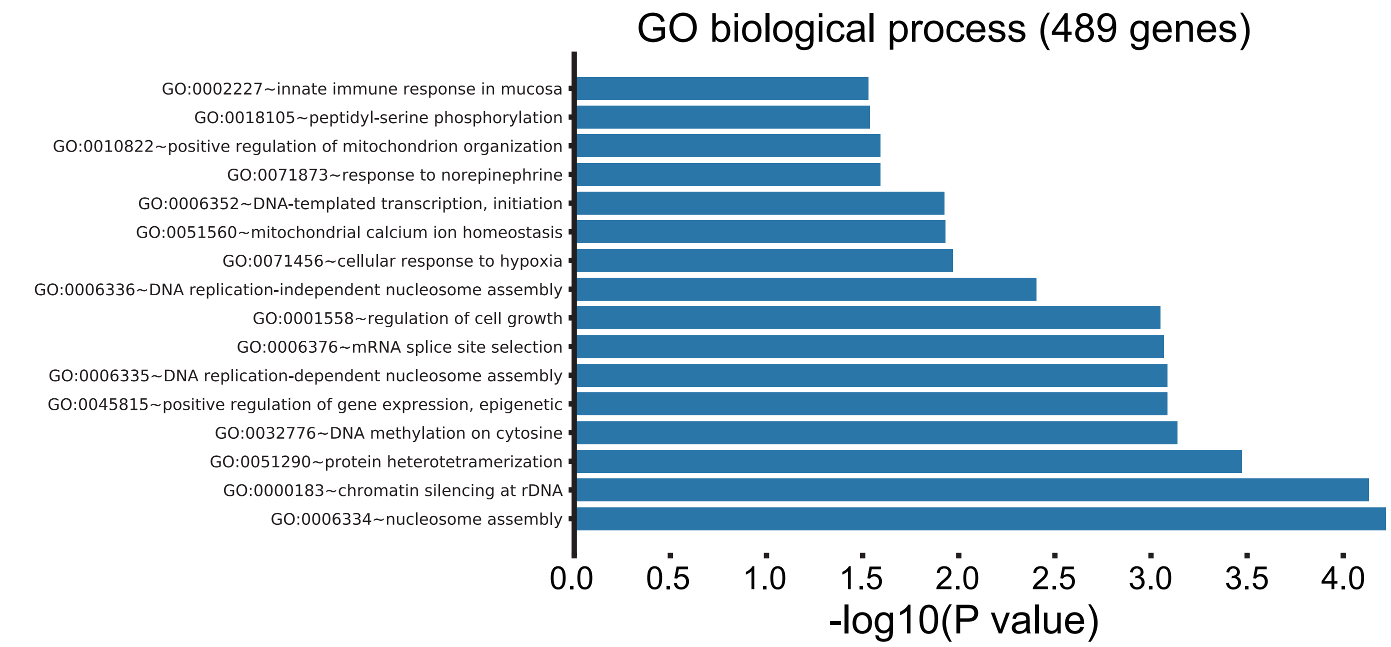


b

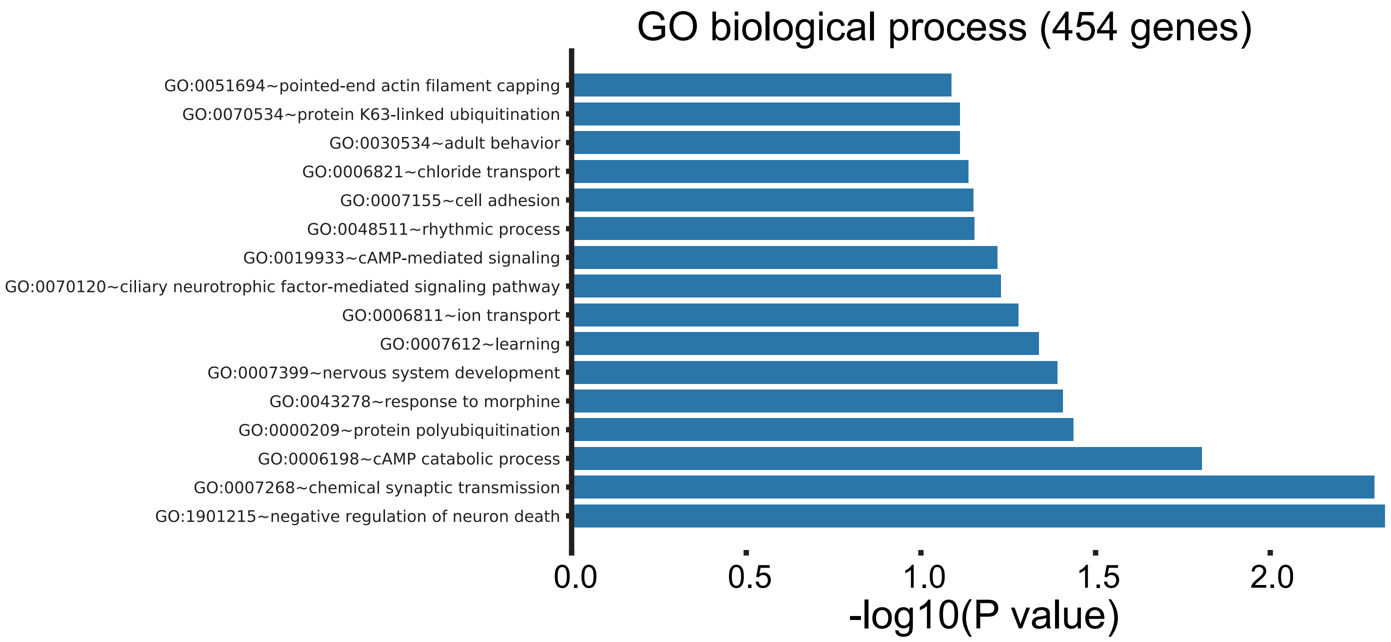


c

| Chip-seq peaks | 2i specific single-cell TAD boundaries(n=346) | serum specific single-cell TAD boundaries(n=351) | Binomial test (two-tailed) |
|---|---|---|---|
| ZC3H11A | 77 | 95 | pvalue=0.195 |
| CHD2 | 33 | 34 | pvalue=0.903 |
| MAFK | 65 | 64 | pvalue=0.930 |
| H3K36me3 | 423 | 546 | pvalue=0.000 |
| H3K9me3 | 415 | 341 | pvalue=0.004 |
| CTCF | 181 | 237 | pvalue=0.008 |
| H3K4me3 | 118 | 165 | pvalue=0.007 |
| HCFC1 | 45 | 74 | pvalue=0.008 |
| ZNF384 | 147 | 165 | pvalue=0.365 |
| H3K4me1 | 32 | 15 | pvalue=0.013 |
| H3K9ac | 269 | 345 | pvalue=0.004 |
| H3K27ac | 195 | 206 | pvalue=0.653 |

d

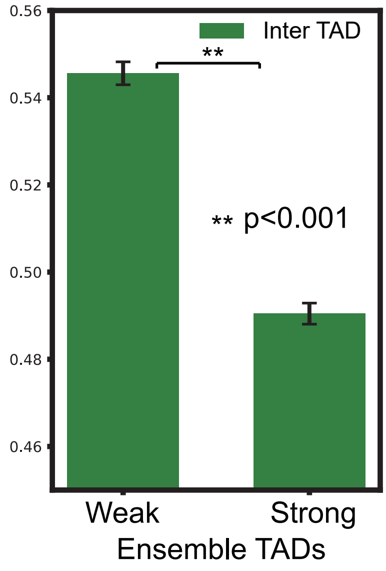


e

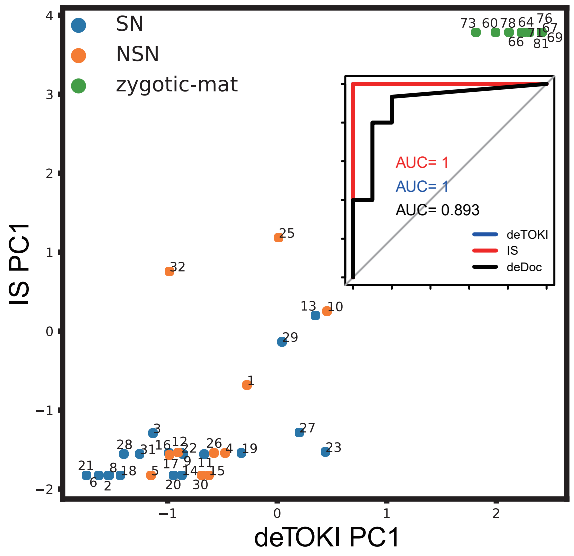


Figure S8
