## supplemental text for "DeTOKI identifies and characterizes the dynamics of chromatin topologically associating domains in a single cell"

**The performance of deTOKI in bulk Hi-C and simulated Hi-C**

We validate the efficiency of deTOKI in Dixon's Hi-C data. It shows that TAD boundaries locate on the boundaries of block pattern in contact matrix (Supplementary Fig. 1a). Aggregate analysis of deTOKI-predicted TADs also reveals the enrichment of functional signaling on boundaries, including CTCF, H3K4me3, and H3K36me3. (Methods; Supplementary Fig. 2a-c). To cross-validate the accuracy of deTOKI's boundary, we compare the similarity between it and the results of other TAD algorithms, namely Insulation Score (IS) and deDoc (Methods). deTOKI has high similarity IS, a widespread method used in recent years (Supplementary Fig.2d). The deTOKI also performs well in simulated Hi-C data based on CTCF motif. It is well known that CTCF motif often shows convergent orientation at chromatin interactions, as well as bias of orientation upstream or downstream of TAD boundary. Therefore, we define "motif insulation strength" to simulate a Hi-C contact matrix and perform deTOKI (Methods). As expected, the detected domains based on CTCF motif are very similar to TADs detected from Hi-C data (Supplementary Fig.2e).

To simulate ensemble Hi-C data based on CTCF motif, we define motif insulation strength between $bin_i$ and $bin_j$, as follows:

$$Insulation_{ij} = max_{i\le k\le j}\left(\sum_{i\le x<k} BM_x + \sum_{k\le x<j} FM_x\right),$$

where $BM_x$, $FM_x$ denote the backward and forward motif number in $bin_x$, respectively. Then we perform Poisson regression on existing Hi-C data as follows to simulate the Hi-C data:

$$Contact\ \ number_{ij} \sim 1 + abs(j-i) + Insulation_{ij}.$$

**Supplementary Figure legends**

**Supplementary Fig.1 A** The predicted TADs under different "k" (NMF times) in four examples, including contact matrices of ensemble Hi-C data (in Dixon et al.) and single-cell Hi-C data (in Tan et al.). Predicted TADs are shown in sawtooth. **B** The predicted TADs under different resolution in two examples, including contact matrices of ensemble Hi-C data (in Dixon et al.) and single-cell Hi-C data (in Tan et al.). Predicted TADs are shown in sawtooth. **C** The left and right scatter plots represent running time of deTOKI using 1 core or 16 cores, respectively. Each point represents an intra-chromosome Hi-C contact matrix from oocytes ensemble Hi-C data (in Flyamer et al.).

**Supplementary Fig.2** deTOKI can accurately detect TADs in ensemble Hi-C data. **A-C** The plots represent the expectancy of ChIP-seq peaks with CTCF, H3K4me3, and H3k36me3 on the predicted ensemble TAD boundaries (chr1-22), respectively. The *y*-axis represents the mean number of peaks per bin with the same distance to the predicted TAD boundaries. The shadow represents 95% confidence interval as calculated by bootstrap. The p value is resulted from permutation test on enrichment of ChIP-seq peaks on TAD boundaries. **D** The radar plot shows the similarities between the TADs predicted by the different algorithms. Each spoke represents a comparison of AMIs between a reference algorithm (indicated as a colored square) and each of the other algorithms. Abbreviations: IS (Insulation Score), DD (deDoc), MT (MrTADFinder), HM (HOMER), AT (Armatus), HS (HiCseg), TK (deTOKI). **E** An example of simulated data based on CTCF motifs. Heatmap of the Hi-C contact matrix from bulk Hi-C data, and simulated data are the upper part and lower part, respectively. Predicted TADs and CTCF domains are shown in blue sawtooth and green sawtooth, respectively.

**Supplementary Fig.3** Comparison of TAD predictors in simulated single-cell Hi-C data. **A** The differences of TADs, as inferred by AMI and WS, between raw data and down-sampled data in different chromosomes. **B** The (log2) change on number of predicted TADs in different chromosomes on 20kb bin-size and 80kb bin-size. **C** The similarity and differences of TADs, as inferred by each index, between raw data and down-sampled data in different chromosomes on 20kb bin-size and 80kb bin-size. **D** The genome-wide distribution of ChIP-seq peaks of CTCF, H3K4me3 and H3K36me3 flanking the predicted TAD boundaries**,** respectively. The shadow represents 95% confidence interval as calculated by bootstrap. The *y*-axis represents the mean number of peaks per bin with the same distance to the predicted TAD boundaries (MNPPB). The enrichment p values are calculated by permutation test (n=10000). **E** From left to right, the normalized Hi-C contact matrix of chr18:10-15Mb for GM12878 ensemble Hi-C from Rao's data[18], an ensemble of 100 modeled 3D structures of this region, and the 3D structure modeled from the simulated ensemble Hi-C from model #100. Each dot in the right panel represents a particle 10kb long, and the dots with same color belong to the same predicted ensemble TAD. **F** The differences of predicted single-cell TADs between different thresholds and predictors on chr18:50-55Mb. **G** The cumulative distribution function of distance between bin pairs in the representative example (model#1). **H** The similarities and differences of predicted single-cell TADs between different thresholds and predictors on chr18:10-15Mb. *: P<0.05, **: P<0.001, NS: not significant, two-sided Mann-Whitney U test.

**Supplementary Fig.4** deTOKI performs well in real single-cell Hi-C data. **A** Radar plots on the left and right panel show Modularity Index and Structure Entropy of predicted TADs by each software program on chr1 of 30 oocytes and 10 zygotes-mat (in Flyamer et al.) and on chr1 of 150 mESCs (in Li et al.), respectively. **B-C** The probability mass function of length of predicted TADs by deTOKI and IS in single-cell Hi-C data (PBMC cell#14 chr1) and its down-sampled half data. **D** An example of single-cell data (in Tan et al.) and its down-sampled data at half level. Heatmap of the Hi-C contact matrix from single-cell Hi-C data; down-sampled data are the upper and lower panels, respectively. Predicted TADs in each data are shown in sawtooth. **E** The similarities between predicted TADs in several single-cell Hi-C data and their down-sampled data at half level by IS and deTOKI. **F** The probability mass function of TAD length predicted by deTOKI and IS in chr1-22 (in Tan et al.). **G** The mean contact coverage (in Tan et al.) on mini TADs predicted by IS and other TADs in chr1-22. *: P<0.05, **: P<0.001, NS: not significant, two-sided Mann-Whitney U test.

**Supplementary Fig.5** TAD structure is highly dynamic at the single-cell level. **A** The cell-to-cell and cell-to-ensemble similarity of deTOKI-predicted TADs. The single-cell data was from 30 oocytes (in Flyamer et al.), compared to the ensemble in mESC and oocyte. **B** The cell-to-cell and cell-to-ensemble similarity of deTOKI-predicted TADs. The single- cell data was from 150 mESCs (in Li et al.), compared to the ensemble in mESC and oocyte. **C** The diagram of four types of TAD changes in a single cell. **D** Distribution of different types of ensemble TADs in chr1 (in Tan et al.). **E** Example of predicted single-cell and ensemble TADs. The type of ensemble TAD is marked in color. *: P<0.05, **: P<0.001, NS: not significant, two-sided Wilcoxon rank-sum test.

**Supplementary Fig.6** The ensemble TAD boundaries were not purely randomly distributed in single cells. **A** Number of cells in which the ensemble TAD boundary is also a TAD boundary. The statistic is shown for four types of ensemble TAD boundaries. The p value was calculated by two-sided Wilcoxon rank-sum test. **B** and **C** GO analysis of genes on the over- and under-represented boundaries, respectively.

**Supplementary Fig.7** The scSBs may not fully result from stochastic fluctuation. **A** The distribution of histone marks flanking the IS- or deDoc-predicted single-cell boundaries is shown, respectively**. B** The distribution of histone marks flanking the IS- or deDoc- predicted ensemble boundaries are shown, respectively**.** The *y*-axis of panel **(A-B)** represents the mean number of peaks per bin with the same distance to the predicted TAD boundaries normalized by average in whole genome (MNPPB). The shadow represents 95% confidence interval, as calculated by bootstrap. **C** The enrichment p values and consensus p values (Methods) of each histone mark on single cell TAD boundaries and ensemble TAD boundaries predicted by deTOKI. **D** enrichment p values and consensus p values (Methods) of each histone mark on scSBs-m and scSBs-1,2. **E** The average number of ChIP-seq peaks in scSBs. **F** The distance to the nearest ensemble boundaries of scSBs in each class. **G** A logistic regression model to classify scSB-1,2 and scSB-m based on 12 Chip-seq peaks. Factors with a positive coefficient have a direct effect on scSB-m. Only the significant factors are displayed. *: P<0.05, **: P<0.001, NS: not significant, two-sided Wilcoxon rank-sum test.

**Supplementary Fig.8** TAD structure carries the information for the cell identity.

**A-B** GO analysis of genes on the serum-specific single-cell TAD boundaries and genes on the 2i-specific single-cell TAD boundaries, respectively, in Li's dataset. **C** The number of ChIP-seq peaks on two types of single-cell TAD boundaries. **D** The PCC of DNA methylation rate in bin pairs cross ensemble TAD boundaries which have a weak insulation score, and bin pairs cross ensemble TAD boundaries which have strong insulation score. **E** The classification of single cells based on predicted TAD boundaries in Flyamer's datasets. The x- and y-axis represent the PC1 calculated by deTOKI and IS, respectively. The embedded plots show the AUC of classification by each program. *: $P<0.05$, **: $P<0.001$, Fisher's z-test.

**Supplementary Table 1** The statistics of predicted TADs using three methods with Tan's data.
The statistics includes number and length of predicted TADs using IS, deDoc and deTOKI. Each row represents a cell on Tan's data. GM12878#8 was eliminated because it doesn't have cleaned data on GEO.

**Supplementary Table 2** The statistics of predicted TADs using three methods with Flyamer's data.
The statistics includes number and length of predicted TADs using IS, deDoc and deTOKI. Each row represents a cell on Flyamer's data. The names consist with sample names on GEO.

**Supplementary Table 3** The statistics of predicted TADs using three methods with Li's data.
The statistics includes number and length of predicted TADs using IS, deDoc and deTOKI. Each row represents a cell on Li's data.

**Supplementary Table 4** The description of used Hi-C and single-cell Hi-C data. The description includes cell type, resolution of contact matrix, contacts number, sparsity, dynamic range of contacts and availability of data. The calculation method of sparsity and dynamic range is explained at the bottom of the table.
